# *Drosophila suzukii* oxidative stress response involves *Jheh* gene cluster but not transposable elements

**DOI:** 10.1101/2020.04.27.063297

**Authors:** Pierre Marin, Angelo Jacquet, Hélène Henri, Patricia Gibert, Cristina Vieira

## Abstract

The study of the mechanisms involved in adaptation remains a timely issue, particularly in the context of global changes. To better understand these mechanisms of rapid adaptation, invasive species are a good model because they are subjected to new and/or different environmental factors. Using different lines of different geographical origin of the invasive pest *Drosophila suzukii*, we characterized the phenotypic response to oxidative stress. Subsequently, we tested the involvement of the *Jheh* gene cluster in this response and the possible role of transposable elements. We show that the resistance to oxidative stress of the lines appears to be related to their invasive status and we confirm the role of the *Jheh* gene cluster in this response. We have not identified any transposable elements in this gene region that could influence the expression of the gene.

**Summary statement:** The responses to oxidative stress of the invasive species, Drosophila suzukii, show variability between genotypes related to their invasion status. The genes of the juvenile hormone epoxide hydrolase cluster are involved in this response.

## INTRODUCTION

The rapid spread of invasive species in a huge spectrum of environments relies on multiple factors, from genetics to phenotypic plasticity, probably including fine molecular mechanism such as hormonal production or epigenetic gene regulation (Beldade *et al.*, 2011; Marin *et al.*, 2019; Stapley *et al.*, 2015). Phenotypic plasticity, i.e., the ability of a genotype to express different phenotypes in different environments (Ghalambor *et al.*, 2015) has been proposed as one of the most promising explanations for invasive success, particularly in the case of founder population depleted of genetic variation (Estoup *et al.*, 2016; Marin *et al.*, 2019). Among deleterious environments that can be encountered by invasive species, oxidative stress caused by phytosanitary products is one of them. The invasive pest, *Drosophila suzukii,* is a good model to investigate the adaptive process during invasion (Gibert *et al.*, 2016). This species which belong to the group of the fruit fly *D. melanogaster*, originally comes from Asia and was detected simultaneously both in North America (U.S.A) and in Europe in 2008. North America was invaded by native Japan populations derived from Hawaii. In Europe, several introductions were detected from U.S.A and from China (Fraimout *et al.*, 2017). Currently, *D. suzukii* is present in both North and South America, in Europe from the south (Spain) to the East (Poland, Ukraine) and it has also been observed in Russia (CABI, 2020; Lavrinienko *et al.*, 2017).

Characterization of the phenotypic and molecular responses of *D. suzukii* to changing environmental conditions may provide information to the mechanisms involved in the ability of invasive species to cope with environmental variation. Paraquat (N,N′-dimethyl-4,4′-bipyridinium dichloride) is one of the most widely used herbicide in the world leading to the production of ROS (reactive oxygen species) (Tsai, 2018). Oxidative stress due to the use of paraquat in the field has also been used in the laboratory as a good proxy for studying stress resistance (Bus J S and Gibson J E, 1984; Rzezniczak *et al.*, 2011). Paraquat was banned since 2007 in Europe but is still used in many other regions like in U.S.A or Japan. Paraquat exposition is known to induce a reduction in the lifespan associated with changes in gene expression (Finkel and Holbrook, 2000; Liguori *et al.*, 2018; Vermeulen *et al.*, 2005). One of the candidate genes involved in paraquat resistance is the cluster of *Jheh* (*Juvenile hormone epoxide hydrolase*) genes, which are not only involved in the lifespan but also in response to the oxidative environment (Flatt and Kawecki, 2007; Guio *et al.*, 2014). Moreover in *D. melanogaster*, an insertion of a transposable element (TE) *Bari-Jheh,* near the cluster of the *Jheh* genes has been described as driving an increase of resistance in presence of paraquat (Guio *et al.*, 2014).

Using several strains of *D. suzukii*, we measured responses to oxidative stress at the phenotypic and molecular level. We made the hypothesis that different genetic backgrounds from native and invasive populations will have different responses to oxidative stress and that the *Jheh* cluster may be involved on it. Due to the over-representation of TEs in the genome of *D. suzukii* (33% of the repeated elements, (Sessegolo *et al.*, 2016)), compared to other *Drosophila* species, we looked for the presence of TEs in this region in the different lines. We monitored lifespan after paraquat exposure and measured the expression of three genes of *Jheh* cluster *Jheh-1*, *Jheh-2* and *Jheh-3* in six isofemale lines, four from the invasive regions, North America (Watsonville and Dayton) and France (Paris and Montpellier) and two from the native area, Japan (Sapporo and Tokyo). We evaluated the genetic diversity within and between lines by sequencing introns of the *Jheh* genes, searched for TEs and for transcription factor binding sites (TFBS). Our results suggest a strong effect of the genotype on the resistance to stress and changes in *Jheh* expression levels, with no link with TEs.

## MATERIAL AND METHODS

### *Drosophila suzukii* lines and rearing conditions

*D. suzukii* lines were sampled in 2014 from one native country (Japan: Sapporo and Tokyo) and 2 invaded areas (USA: Watsonville and Dayton and France: Montpellier and Paris). Field-inseminated females were isolated to establish half-sib families called isofemale lines commonly used to investigate Drosophila natural populations (David *et al.*, 2005). Flies were reared in modified medium (drosophila agar type, ref.66-103, Apex™,9 g.L^−1^; cornmeal 33 g.L^−1^; yeast, dried yeast, ref.75570, LYNSIDE^Ⓡ^ 17 g.L^−1^; industrial sugar 50 g.L^−1^; nipagin, Tegosept, ref.20-258, Apex™ 4 g.L^−1^; 96% ethanol 40 ml.L^−1^; distilled water 1 L) from Dalton *et al.*, (2011), in a humidified, temperature-controlled incubator at 22.5°C, 70 % of relative humidity and a 16:8 LD cycle. The recipe of the modified medium was to bring to boil agar, cornmeal, yeast extract and sugar in distilled water. Then wait out of the fire about 10 minutes until the mixture cooled to 53°C before adding diluted nipagin in 96% ethanol. Medium is then poured in vials and cooled at room temperature before to be stored at 4°C. All the experiments were made with 4 to 7 days old flies.

### Oxidative stress resistance experiments

We used paraquat (methyl viologen dichloride hydrate, ref. 75365-73-0, Sigma-Aldrich^Ⓡ^) to mimic oxidative stress. Oxidative stress was assessed by adding paraquat directly in the medium (10 mM) before the cooling step and below 53°C. The control experiment was made with the same medium but without paraquat. We used one isofemale line per locality (total of six) named Montpellier (France), Paris (France), Sapporo (Japan), Tokyo (Japan), Dayton (U.S.A.) and Watsonville (U.S.A.). We made three replicates per line and per sex, with ten flies per replicate. Survival was monitored every 24h. Flies were transferred into new vials every three to four days to limit microbial development.

### RT-qPCR analysis of *Jheh* genes

We quantified the expression of the three *Jheh* genes (*Jheh-1, Jheh-2* and *Jheh-3*) by RT-qPCR after induction of oxidative stress and in control condition. Adult males and females were exposed during 24h to medium culture with 20 mM of paraquat. After 24h the flies were immediately dissected in PBS 1X solution (Gibco Thermo-Fisher) in order to extract carcasses for both sexes and eliminate germline tissues. We made three replicates per sex and treatment and used four flies per replicate.

RNA extraction was made using Direct-zol™-96 RNA Kits (Zymo Research), following the manufacturer recommendations and RNA was treated with DNAse. cDNA were obtained from 0.5 μg of RNA using SuperScript™ IV VILO™ Master Mix (Invitrogen). RT negative control was made with RNA but without the reverse transcriptase to control for genomic DNA contamination. cDNA were stored at −80°C before the quantification step. Gene expression was then quantified by quantitative PCR and *Rp49* was used as housekeeping gene. Primers were designed using the *D. suzukii* referenced genome (Table S1, (Chiu *et al.*, 2013)). Their efficiency was between 91.1% to 97.2% (*RP49*: 91.6 %, *Jheh-1*: 97.2%, *Jheh-2*: 95.2%, *Jheh-3*: 91.1%). 2 μl of the cDNA sample were supplemented with 5 μL of SsoADV Universal SYBR Green Supermix (BioRad) mix 2X, 0.3 μl of each primer (10μM) and 2.4 μl of pure water. We made technical duplicates for each sample. PCR reactions were made in a BioRAd CFX-96 with a program consisted of an initial activation of 95°C for 10 minutes and then 40 cycles each comprising 15 seconds at 95°C, 10 seconds at 60°C and 72°C.

### Genetic diversity of isofemale lines

We sequenced intronic regions of *Jheh* gene cluster of the six lines used in this study. DNA was extracted individually from 10 females per line with the 96-Well Plate Animal Genomic DNA Miniprep kit (ref. BS437, Biobasic) following the manufacturer instructions. Primers were designed to flank the intronic regions for the three *Jheh* genes (Table S1) and Phusion high fidelity DNA Polymerase (2 U/μL) (F-530XL Thermofisher Scientific) was used to amplify sequences. The same PCR program was used for all primers pairs: 98°C for 10 minutes, followed by 40 cycles composed of 30 seconds at 98°C, 1 minute at 56°C and 20 seconds at 72°C and a final elongation step for 1 minute at 72°C. The sequencing of the two strands was done directly from the PCR product by BIOFIDAL sequencing company (Vaulx en Velin, France). Sequences were manually curated with CLC Main Workbench 8 software (Qiagen) before being aligneed with the Muscle program implemented in the workbench to generate haplotypes by line for each intron. MEGA X software was used to calculate pairwise comparison and nucleotide diversity using p-method option (Table S2–S3) (Kumar *et al.*, 2018).

### Detection of Transposable elements and transcription factor binding sites

We sequenced the intergenic regions of the *Jheh* gene cluster, plus the 5’ and 3’ regions of the cluster (Table S1). DNA was extracted from one female per population as described above. Classical PCR method was used with the following program, 10 minutes at 95°C followed by several cycles composed of 30 seconds at 95°C, 30 seconds at 63°C, 3 minutes at 72°C and a final elongation of 15 minutes at 72°C. The number of cycles was25 for the region before *Jheh-1* and between *Jheh-1* and *Jheh-2*, 35 cycles for the region between *Jheh-2* and *Jheh-3* and 30 cycles after *Jheh-3*. We identified TEs in the intergenic regions by a blast against a homemade data base of the TE sequences from the *D. suzukii* reference genome (Paris *et al.*, 2020, Mérel *et al.*, *in prep.*).

For TFBS (Table 1), we used conSite website to screen all TFBS from insect in our sequences (Sandelin *et al.*, 2004). To complete our analysis, we used the TFBS obtained from Villanueva-Cañas *et al.* (2019) and we extracted PFM (position frequency matrix) of the 14 TFBS from the JASPAR2018 database (v.1.1.1) (Parcy *et al.*, 2017). Then, we used TFBSTools (v.1.22.0) package from R software (v. 3.6.0) to convert in PWM (position weight matrix), and then search on the 6 lines and the reference genome of *D. suzukii (Paris et al., 2020; R Core Team, 2019; Tan and Lenhard, 2016).*

**Table 1.**
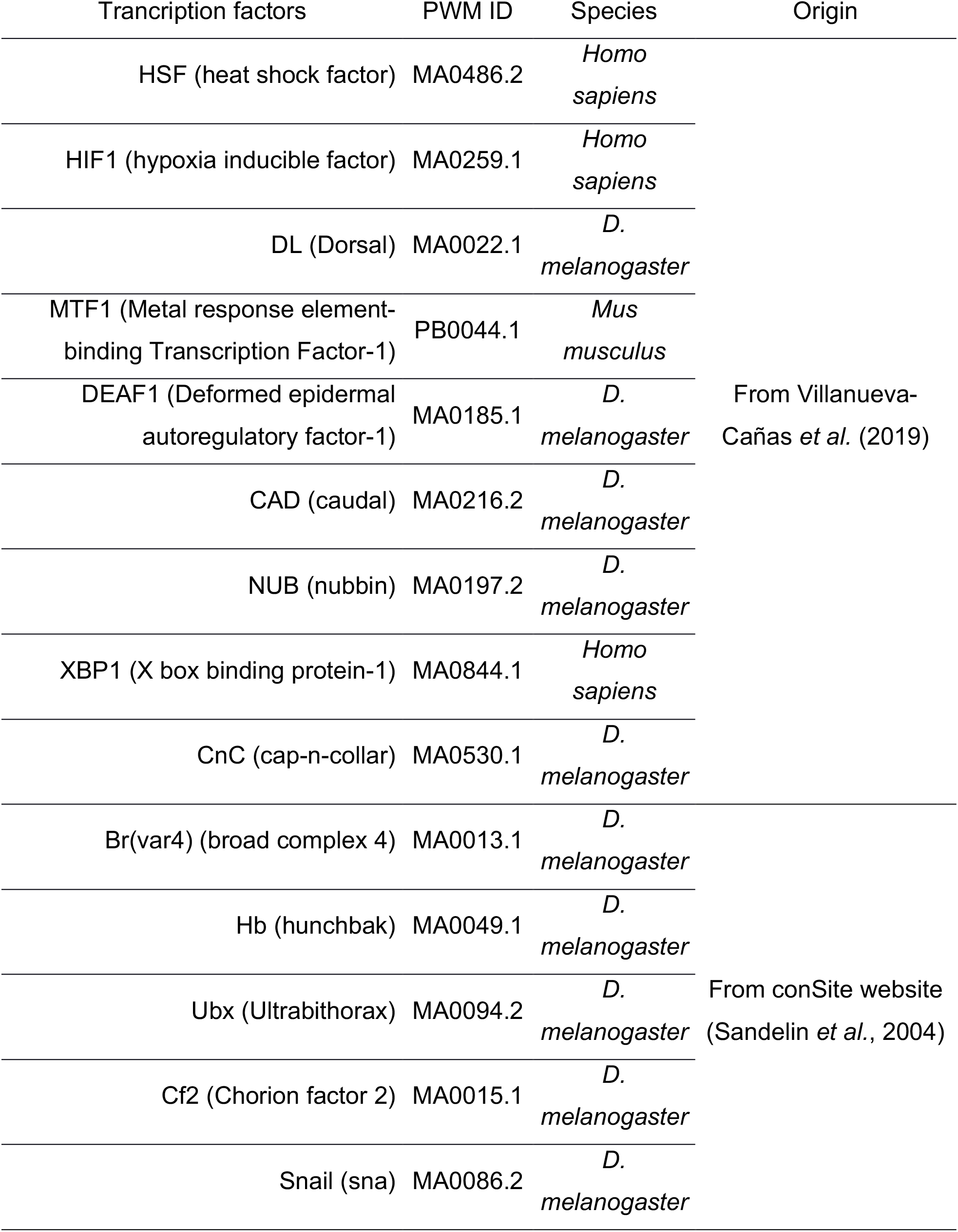
Transcription factor analyzed with the PWM matrix ID from JASPAR2018. Mainly matrix come from *D. melanogaster* model, but several as HSF, HIF1 and XBP1 come from human, while MTF1 come from mice.

### Statistical analysis

### Survival monitoring

Survival data were analyzed using a linear mixed model with lmer function from lme4 provided on R (v. 3.6.0) after a log transformation, confirmation of normality and homoscedasticity (Bates *et al.*, 2015). This model was chosen after log-likelihood comparison between models (linear model with raw or log transformed data, survival model with a Weibull distribution).

We analyzed sexes separately to limit interaction terms, and focused on the effect of the treatment, the lines and their interaction. Biological replicates were added as random effect and we plotted exponential of the values and associated confidence interval on the Fig. S1. Those effects can be interpreted as multiplicative effect on the mean lifespan compared to the reference chosen here as the non-exposed group from Sapporo (*e.g.* the Sapporo reference is centered on 1 and the effect of paraquat 0.18 involves a survival time under paraquat for Sapporo of 0.18 or 18% of the survival time without paraquat).

### qPCR analysis

RT-qPCR raw data were analyzed using R and EasyqpcR library (1.21.0) for the quantification and normalization with *RP49* (Sylvain, 2012). Data were analyzed separately for the three genes (*Jheh-1 -2* and *-3*) and sex using a linear model (ANOVA2, Table S4) after log transformation to validate homoscedasticity and normality. Pairwise comparisons were made using a Tuckey test.

## RESULTS

### *D. suzukii* wild type lines have significant differences in life span

To investigate the influence of the genotypes from different geographical origins on the lifespan, we compared the invasive and native *D. suzukii* lines in control condition (Fig. 1A, Fig. S1 & Table 2). The lifespan ranges from 31 to 55 days for females and from 25 to 45 days for males. For females we observed a strong genotype effect related to geographic location: the genotypes that lived the longest were those of Dayton and Paris (about 1.88-1.96 times more than Sapporo, Fig. S1). Sapporo, Tokyo, and Watsonville were not significantly different and with the lowest lifespan. For males, the four invasive genotypes from Europe and U.S.A had a higher lifespan than Sapporo. Tokyo was similar to Sapporo.

**Table 2.** Mean (±SD) survival time (days) for male and female lines of D. suzukii under control and paraquat conditions. Sensitivity represents the exponential of the interaction values in the model (i.e., the difference in slope between the Sapporo reference and the other genotypes, see Figure S1). * indicate a significant difference with the reference (p-value < 0.05).

| Females |  |  |  | Males |  |  |
| --- | --- | --- | --- | --- | --- | --- |
| Lines | Control | Paraquat | sensitivity | Control | Paraquat | sensitivity |
| Sapporo (Japan) | 31 ±13.3 | 5.4 ±2.3 | 1.0 | 25.4 ±5.9 | 6.2 ±2.1 | 1.0 |
| Tokyo (Japan) | 27.1 ±9.1 | 5.1 ±1.9 | 1.06 | 29.2 ±11.5 | 7.7 ±4.9 | 1.02 |
| Dayton (U.S.A) | 53 ±8.4 | 9.5 ±3 | 0.98 | 42.8 ±16.3 | 11 ±4.1 | 1.23 |
| Watsonville (U.S.A) | 34.9 ±13.8 | 6.6 ±2.3 | 1.13 | 34.5 ±11.2 | 5.5 ±2.1 | 0.68* |
| Paris (France) | 55.4 ±9.4 | 7.2 ±1.8 | 0.72* | 41.9 ±14.8 | 8.1 ±2.8 | 0.9 |
| Montpellier (France) | 37.9 ±8.4 | 5.7 ±2.2 | 0.83 | 44.7 ±12.2 | 7.5 ±3.8 | 0.66* |

**Fig. 1.**
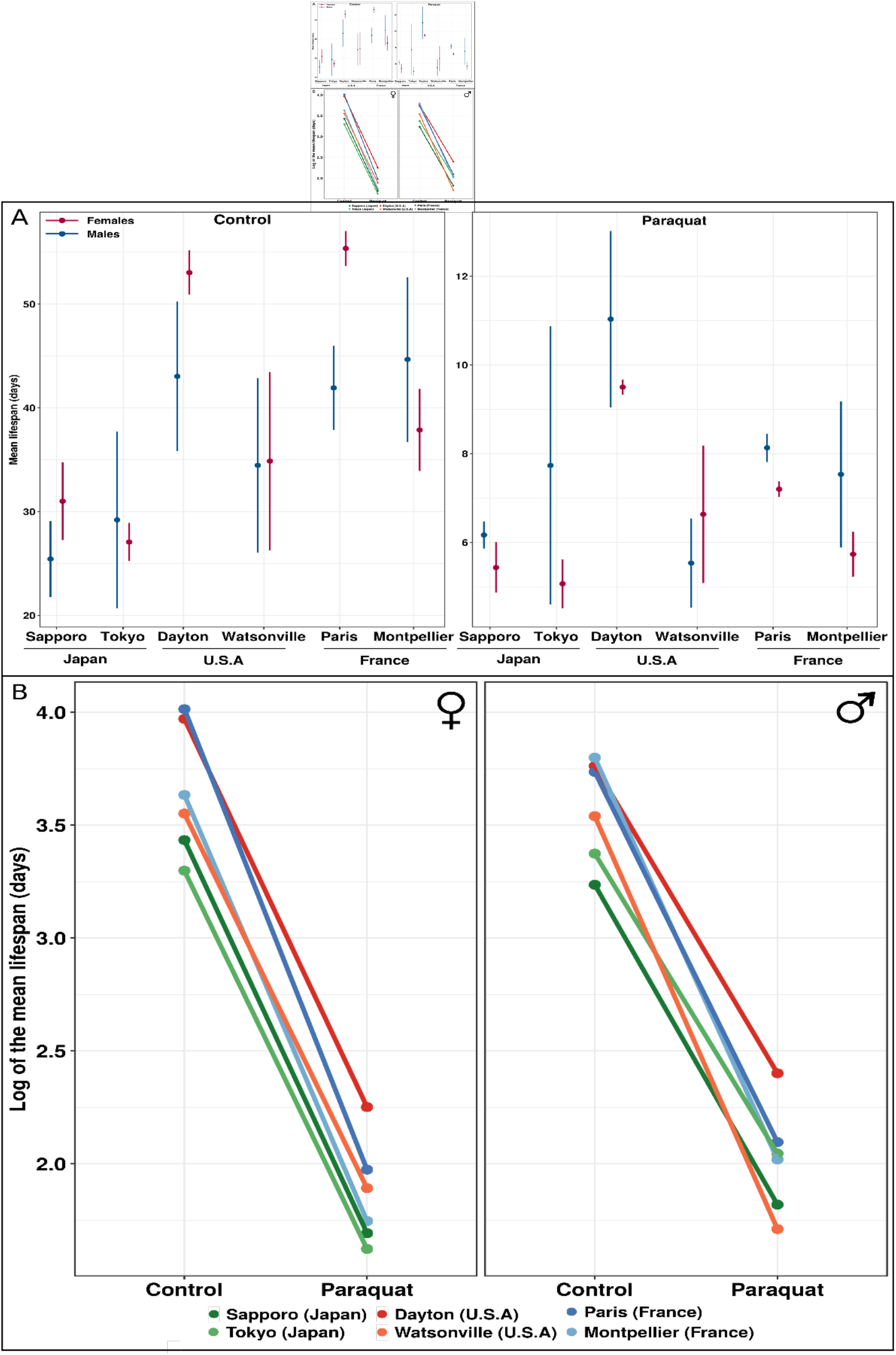
Mean (±SD) of survival time (days) of *D. suzukii* genotypes (A) and reaction norm (log of the mean) (B). (A) mean values under control and paraquat condition for females (pink) and males (blue) with associated SD. (B) reaction norm between treatments (control and paraquat) for both females and males for the six genotypes using log transformed mean value. The slope differences between the curves highlight the difference of sensitivity between genotypes.

As expected, exposure to paraquat reduced life span on average from 82 to 77% for females and males (Fig. 1A, Fig. S1). The two lines with the best paraquat resistance in absolute value were still Dayton and Paris in both sexes (Fig. 1A and Table 2). We then wanted to have an estimate of paraquat sensitivity (i.e., the slope difference Fig.1 B) taking into account the longevity of each line by estimating the value of the interaction coefficients (*i.e*., the slope difference compared to Sapporo) in Table 2 and statistically tested in Fig. S1. Again, the effect was not similar between genotypes and sexes. For females, Paris was the line presenting significantly the highest sensitivity (−0.87, Table 2) with a reduction of 28% of the life span compared to Sapporo (Fig. S1). For males, the reduction in life span was significantly the highest for Montpellier and Watsonville (−0.84 and −0.83) with a reduction from 34 and 32% by comparison with Sapporo. These results reveal a strong genotype-by-environment interaction in the response to oxidative stress and also a sex effect. It is interesting to note that despite the shorter life span of Japanese genotypes, and in particular of Sapporo in the absence of treatment, these genotypes were the most resistant to paraquat exposure, as shown by the lowest ratio of paraquat lifetime to control lifetime (Table 2).

### *Jheh* genes expression changes with the paraquat treatment

To investigate the effect of paraquat-mediated oxidative stress on the gene expression level, we focused on *Jheh* gene cluster described as potentially involved in stress response in insects and mammals (Guio *et al.*, 2014; Oesch *et al.*, 2000). We quantified the level of expression of the *Jheh* genes (*Jheh-1, Jheh-2 and Jheh-3*) in adult males and females flies for the six genotypes described above (Fig. 2).

**Fig. 2.**
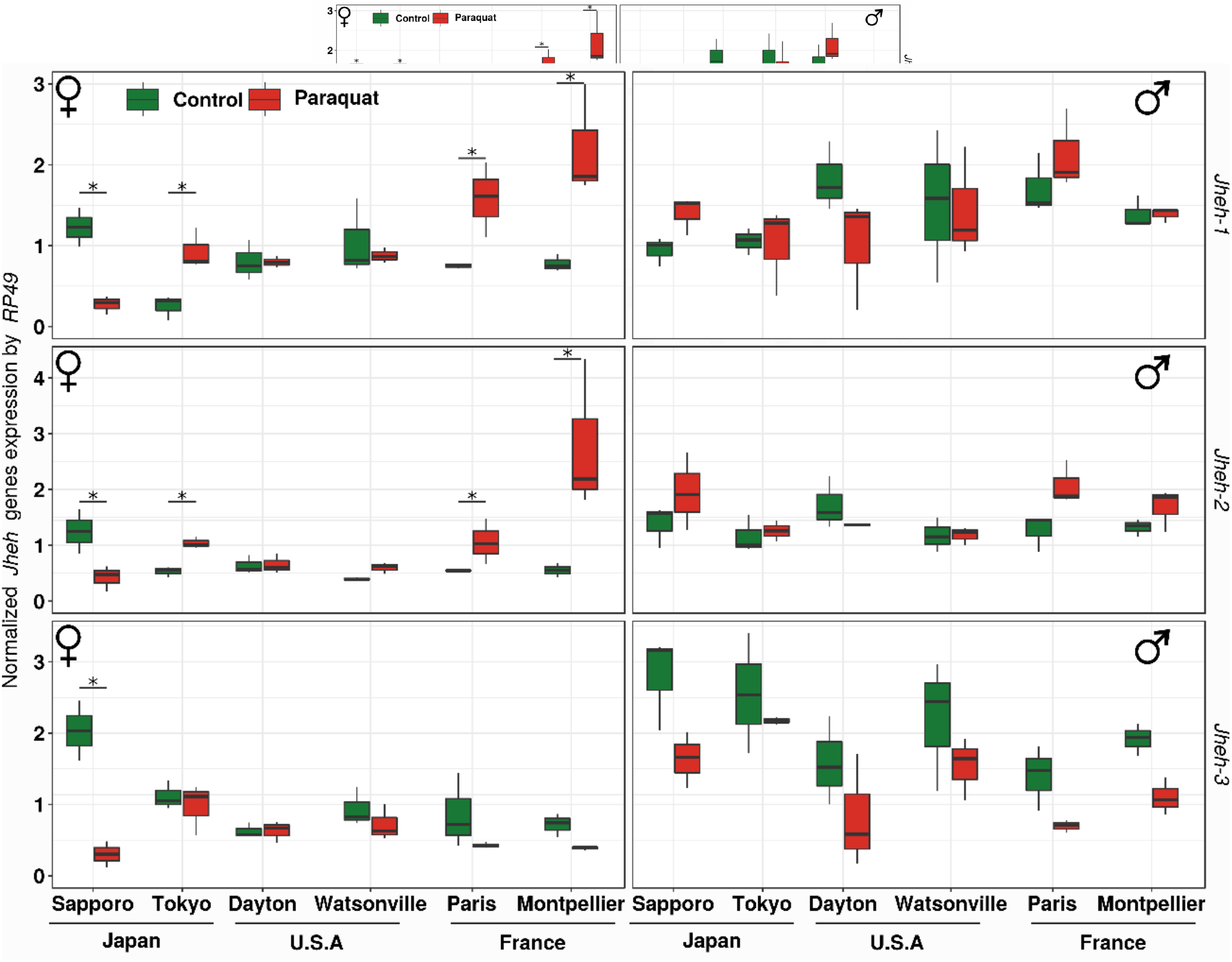
*Jheh* gene expression (*Jheh-1*, *-2*, *-3*) after normalization by *Rp49* in control (green) and paraquat (red) conditions for females on the left and males on the right.

We observed strong differences between males and females. For males, gene expression was not significantly different between control and paraquat treatment for the six genotypes and for the three gene. In females, the effect of paraquat was different according to the gene and the genotype (Table S4). For *Jheh-1* and *Jheh-2*, oxidative stress resulted in a significant increase of gene expression for the two French genotypes and the Tokyo genotype. On the contrary, the Sapporo genotype exhibited a significant reduction of *Jheh-1* expression in presence of paraquat. For *Jheh-3,* we observed a downregulation of the gene expression only for the Sapporo genotype.

### Low Genetic diversity of lines in *Jheh* cluster

To assess the levels of neutral genetic diversity within and between lines, we sequenced intronic regions for *Jheh* genes for each genotype (Fig. 3). As expected, the within-line polymorphism was very low (Table 3, Fig. S2), with the exception of Watsonville with 0.0792 for the first intron of *Jheh-1*. The number of haplotypes was also low (Table S2). The first intron of *Jheh-2* presents the highest levels of diversity, contrasting with the other introns. This corresponds to a residual polymorphism that is still present in the lines despite the laboratory rearing.

**Table 3.** Within-line diversity (pi) for the six genotypes of D. suzukii for the seven sequenced intronic regions. The mean diversity per intron was calculated using the most common sequence of the six genotypes.

|  | *Jheh-1.1* | *Jheh-1.2* | *Jheh-1.3* | *Jheh-2.1* | *Jheh-2.2* | *Jheh-2.3* | *Jheh-3.2* |
| --- | --- | --- | --- | --- | --- | --- | --- |
| **Paris** | 0.0000 | 0.0000 | 0.0000 | 0.0460 | 0.0000 | 0.0000 | 0.0000 |
| **Montpellier** | 0.0000 | 0.0169 | 0.0000 | 0.0502 | 0.0081 | 0.0000 | 0.0339 |
| **Sapporo** | 0.0000 | 0.0000 | 0.0396 | 0.0546 | 0.0000 | 0.0000 | 0.0000 |
| **Tokyo** | 0.0000 | 0.0000 | 0.0000 | 0.0324 | 0.0000 | 0.0144 | 0.0113 |
| **Dayton** | 0.0000 | 0.0000 | 0.0000 | 0.0449 | 0.0000 | 0.0096 | 0.0000 |
| **Watsonville** | 0.0792 | 0.0000 | 0.0198 | 0.0466 | 0.0000 | 0.0000 | 0.0000 |
| **Mean pi**<br>**diversity** | 0.0070 | 0.0080 | 0.0177 | 0.0390 | 0.0214 | 0.0333 | 0.0209 |

**Fig. 3.**
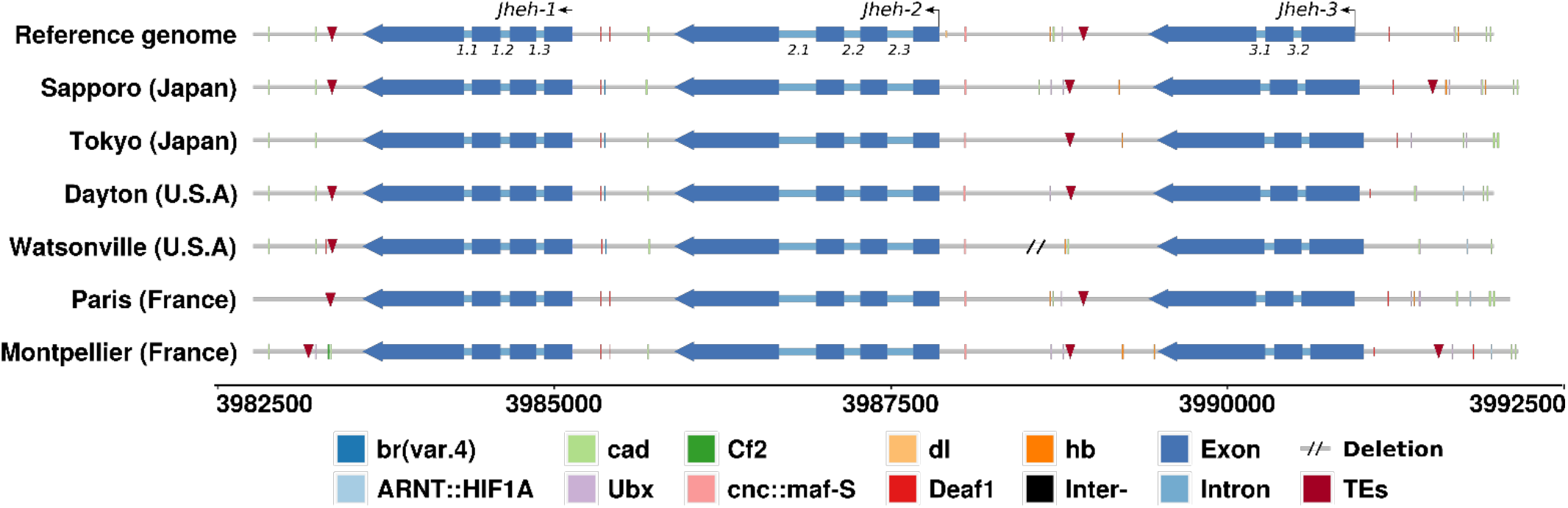
Representation of the Jheh gene cluster on the reference genome and the relative position on the other six genotypes. The blue squares represent the exons and the blue lines the intronic regions. Red triangles represent TE detected in the intergenic regions and TFBS are indicated by vertical lines. The scale is shown below the figure in base pairs.

Depending on the intronic regions we found between two to four haplotypes per genotype (Table S2). We computed the diversity between genotypes (global intronic nucleotide diversity π) using the most common haplotype for each intron, showing that on average these values are very small, with the highest value for *Jheh*-2.1 as mentioned above (Table S3).

### Jheh harbour transcription factor binding sites (TFBS)

Transcription factor binding site (TFBS) are *cis* regulatory sequences that are recognized by transcription factors and modify gene expression. Several TFBS are known to be involved during oxidative response. We detected 9 of the 14 previous identified TFBS in the intergenic regions: HIF1A, br, cad, Cf2, Deaf1, CnC, dl, hb and Ubx (Fig. 3 & 4, Table S5). Comparison of the number of TFBS between genotypes (Fig. 4) revealed several differences but not clear link with the changes in expression observed for the *Jheh* genes. For example, the Sapporo genotype which consistently showed a decrease in the expression of all three genes, did not appear to have a different specific TFBS. The two French genotypes which exhibited systematically an increase of expression after paraquat treatment appeared to have an increased number of putative TFBS. For example, the two French genotypes showed two Deaf1 motives when compared to the other genotypes upstream of TSS of the *Jheh-1* gene. In the case of the *Jheh-2* gene, the French genotypes presented a significant number of TFBS, with for example six TFBS for the Montpellier genotype (2Ubx, 2hb, CnC and cad). In this region, no genotype showed the same pattern of TFBS and it was similar for *Jheh-3*.

**Fig. 4.**
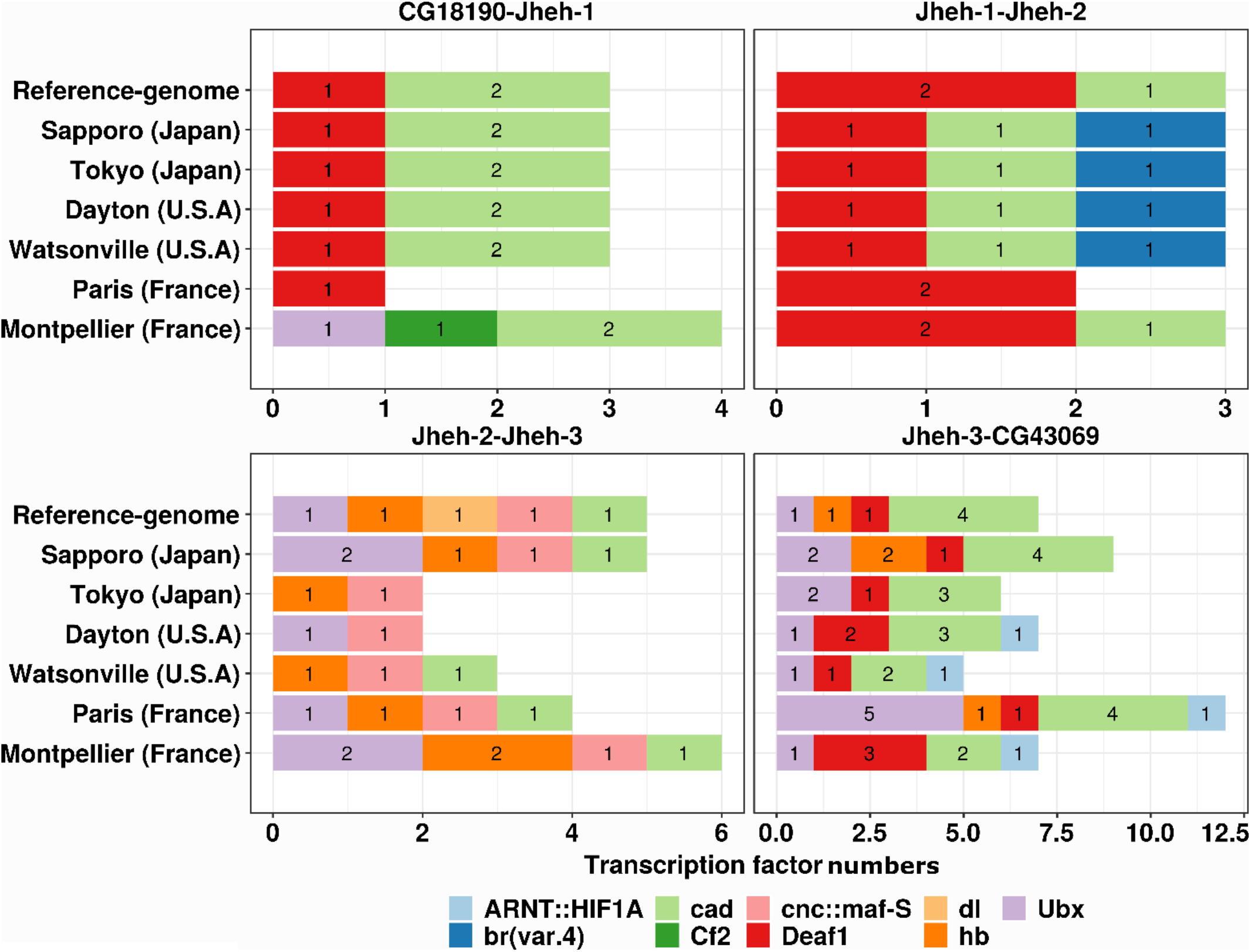
TFBS detected in the four intergenic regions of the Jheh gene cluster in the reference genome and in the six genotypes of *D. suzukii*. Of the 14 TFBS examined, 8 had at least one positive result. The intergenic regions are upstream Jheh-1(CG18190-Jheh-1), between Jheh-1 and -2 (Jheh1-Jheh2), between Jheh2 and -3 (Jheh2-Jheh3) downstream of Jheh-3 (Jheh3-CG43069).

### Transposable elements do not affect *Jheh* gene expression

The presence of TE in the vicinity or within the *Jheh* cluster could impact the gene expression during oxidative stress because they could bring Antioxidant Response Element for transcription factors or by modifying chromatin state (Guio and González, 2015; Guio *et al.*, 2014). Surprisingly, and even if *D. suzukii* harbors more than 30% of TEs, no full insertion was observed in the *Jheh* cluster indicating that we are probably in regions of high recombination. However, we did identify small pieces of TE that are quite conserved between the genotypes but no TFBS were detected in these sequences (Fig. 3, Table 4 & S5). No obvious link seems to exist between gene expression and the presence of TE in the *Jheh* cluster.

**Table 4.** Size differences (bp) between the six genotypes and the reference genome. TE insertions are indicated by their size or abs if they are absent.

|  |  | expected<br>size<br>(reference<br>genome) | Sapporo<br>(Japan) | Tokyo<br>(Japan) | Dayton<br>(U.S.A) | Watsonville<br>(U.S.A) | Paris<br>(France) | Montpellier<br>(France) |
| --- | --- | --- | --- | --- | --- | --- | --- | --- |
| X-Jheh1 | Size | 809 | 0 | -2 | 0 | +1 | 0 | +12 |
|  | TE<br>insertion | 82 | 82 | abs | 82 | 82 | 82 | abs |
| Jheh1/jheh2 | Size | 756 | -17 | -6 | -2 | +2 | +1 | +7 |
|  | TE<br>insertion | abs | abs | abs | abs | abs | abs | abs |
| Jheh2/Jheh3 | Size | 1541 | +31 | -64 | +35 | -297 | -1 | +66 |
|  | TE<br>insertion | 41 | 48 | 49 | 91 | abs | 41 | 49 |
| Jheh3-Y | Size | 1021 | +171 | -8 | +58 | -55 | +29 | +122 |
|  | TE<br>insertion | abs | 58 | abs | abs | abs | abs | 36 |

## DISCUSSION

A growing body of literature suggests that responses to oxidative stress in Drosophila may be mediated by insertions of TEs, that in some cases could affect gene expression or the chromatin structure (Guio *et al.*, 2014). In *D. melanogaster*, the *Jheh* gene cluster has been shown to be involved in the response to paraquat treatment and associated with local adaptation. Guio *et al.* (2014) compared *D. melanogaster* genotypes with and without Bari-Jheh TE insertion, and showed that, (i) TE insertion had a cost in the absence of stress, (ii) TE insertion confer increased survival in the presence of oxidative stress, (iii) TE insertion provides antioxidant response elements (AREs) that contribute to altered gene expression (Guio and González, 2015; Guio *et al.*, 2014). In this study, we analyzed the expression of the *Jheh* gene cluster in several genotypes of *D. suzukii* to test whether the *Jheh* gene cluster is involved in the oxidative stress response and whether TEs could also be associated with alterations in gene expression.

We first measured the life span of flies without treatment. We showed that flies of the Japanese genotypes exhibited the shortest lifespan in both males and females. Surprisingly, these lines showed an increased resistance to oxidative stress. The French lines were more sensitive to paraquat than the American ones, although notable differences were observed between lines from the same continent. The negative association we observed between longevity and paraquat resistance had not been observed in previous work with *D. melanogaster*, in which the opposite association was observed (Finkel and Holbrook, 2000; Liguori *et al.*, 2018). It could be argued that the use of paraquat in Europe has been banned since 2007, which could lead to a loosening of selection on genes related to paraquat resistance, as observed in other organisms (Campos *et al.*, 2014; Shaw, 2000).

We then measured the expression of the *Jheh* genes previously reported to be involved in the oxidative response. Consistent with the literature of *D. melanogaster*, we found sex-specific responses to oxidative stress (Guio *et al.*, 2014; Weber *et al.*, 2012). For *Jheh-1* and *Jheh-2*, we observed a significant effect of genotype and treatment, but only for females, contrary to what was reported in *D. melanogaster*. For *Jheh-3*, treatment and genotype effect were significant for both males and females. These differences in gene expression could not be associated with the presence of TEs insertions, since only partial sequences were present in the intergenic regions. The presence of various TFBS could contribute to the observed differences. We also quantified the polymorphism in our lines, which could be associated with differences in gene expression. We did not observe total homozygosity in the lines but genetic diversity was much lower than what is observed in natural populations of *D. melanogaster*. Lack *et al.* (2016) studied populations from several continents and measured values of nucleotide diversity of up to 0.401 within the population. For inbred DGRP (Drosophila Genetic Reference Panel) lines, the mean intronic diversity was 0.0076 ±0.008, which is close to the values we observed (MacKay *et al.*, 2012). It is therefore unlikely that the residual polymorphism in the *Jheh* gene region can explain the differences in gene expression.

The striking result in our analysis is the similar pattern of changes in the expression of *Jheh-1* and *Jheh-2* in females of European genotypes, with an increase in expression, which was associated with lower resistance to oxidative stress, since these are the most sensitive genotypes. On the contrary, the Sapporo genotype systematically showed a reduction in the expression levels of the three genes, which could also be associated with increased resistance in the presence of paraquat, but this was not observed for the Tokyo genotype.

## Conclusion

In conclusion, our work shows for the first time how various genotypes of *D. suzukii* respond to oxidative stress and suggests that populations found in invaded areas are more sensitive than Japanese populations, specially the French ones. We have also confirmed that the *Jheh* gene cluster is involved in the response to oxidative stress also in *D. suzukii*, independently of the presence of TE in intergenic regions. This work also suggests that the genetic background and probably trans regulatory sequences are involved in gene expression and stress response. Further phenotypic and genomic studies on natural populations are needed to better understand the success of invasive species such as *D. suzukii*.

## Acknowledgements

We thanks to Vincent Mérel who produced the TEs annotation. P. M. produced data, conceived and wrote the manuscript draft. C. V. & P.G. designed experiments, edited the manuscript. A. J. calibrated experimental design. H. H. helped to q-PCR method and analysis.

## List of supplementary figures

**Fig. S1.**
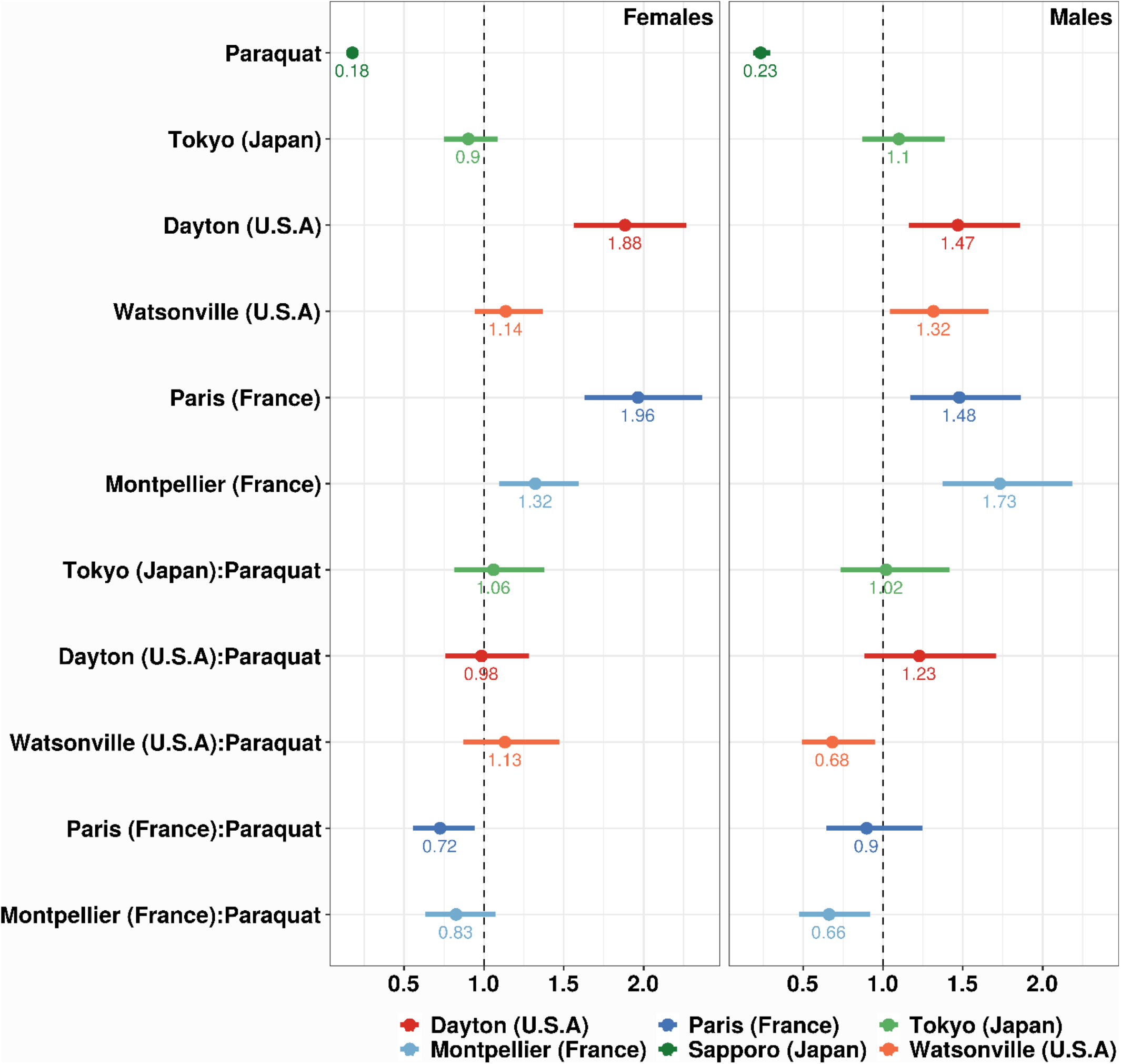
Representation of the parameters estimated by the model for treatment, genotypes and interactions for females (left) and males (right). Values were transformed exponentially to be interpreted as a multiplicator effect. The vertical line corresponds to the reference. Associated p-values are greater than 0.05 when the confidence interval includes the vertical line.

**Fig. S2.**
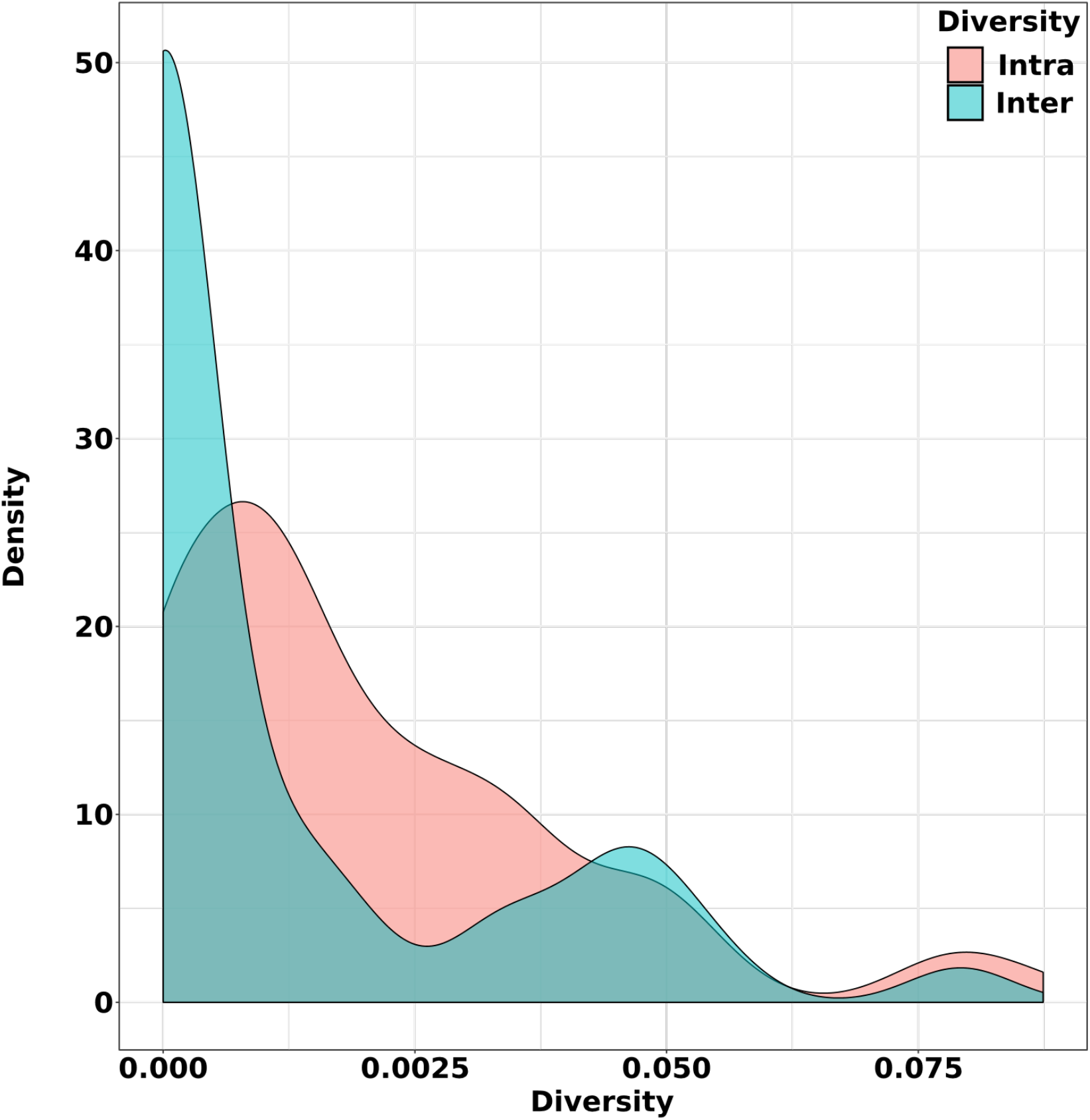
Distribution of the between genotype (blue) and within genotype (pink) genetic diversity using pi values for all intronic regions.

## List of supplementary tables

**Table S1.**
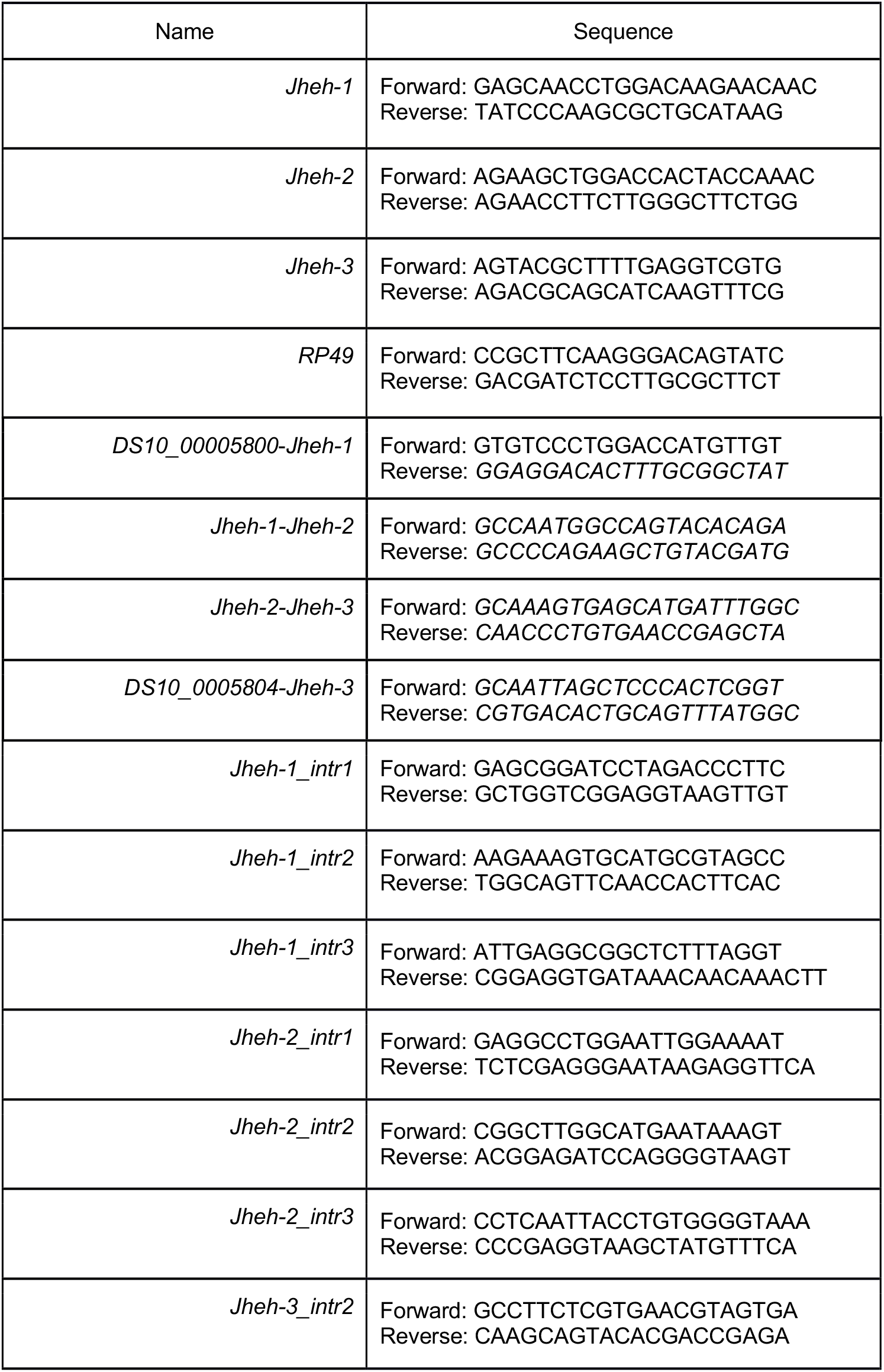
Primers used for the PCR experiments.

**Table S2.** Genotypes of *D. suzukii* with the number of haplotypes obtained. The length of the sequence corresponds to the size of the amplified fragment used to calculate the average diversity per population per intron. The mean diversity was calculated as the average between the most common haplotypes of the 6 lines.

| Lineage |  | Jheh-1 intron 1 |  |  |
| --- | --- | --- | --- | --- |
|  | # of haplotypes | # of flies | Size of the alignment | Mean diversity |
| Paris (France) | 1 | 6 | 97pb | 0.0070 |
| Montpellier (France) | 1 | 10 |  |  |
| Sapporo Japan) | 1 | 6 |  |  |
| Tokyo (Japan) | 1 | 10 |  |  |
| Dayton (U.S.A) | 1 | 10 |  |  |
| Watsonville (U.S.A) | 2 | 7-2 |  |  |
|  |  | Jheh-1 intron 2 |  |  |
|  | N haplotype | N | Alignment | Mean diversity |
| Paris (France) | 1 | 10 | 119pb | 0.0080 |
| Montpellier (France) | 2 | 4-3 |  |  |
| Sapporo Japan) | 1 | 10 |  |  |
| Tokyo (Japan) | 1 | 10 |  |  |
| Dayton (U.S.A) | 1 | 10 |  |  |
| Watsonville (U.S.A) | 1 | 10 |  |  |
|  |  | Jheh-1 intron 3 |  |  |
|  | N haplotype | N | Alignment | Mean diversity |
| Paris (France) | 1 | 10 | 133pb | 0.0177 |
| Montpellier<br>(France) | 1 | 10 |  |  |
| Sapporo Japan) | 2 | 7-2 |  |  |
| Tokyo (Japan) | 1 | 10 |  |  |
| Dayton (U.S.A) | 1 | 10 |  |  |
| Watsonville<br>(U.S.A) | 2 | 5-5 |  |  |
|  |  | Jheh-2 intron 1 |  |  |
|  | N haplotype | N | Alignment | Mean diversity |
| Paris (France) | 2 | 4-2 | 250pb | 0.0390 |
| Montpellier<br>(France) | 2 | 5-2 |  |  |
| Sapporo Japan) | 2 | 4-3 |  |  |
| Tokyo (Japan) | 4 | 2-2-1-1 |  |  |
| Dayton (U.S.A) | 3 | 4-2-1 |  |  |
| Watsonville<br>(U.S.A) | 2 | 5-4 |  |  |
|  |  | Jheh-2 intron 2 |  |  |
|  | N haplotype | N | Alignment | Mean diversity |
| Paris (France) | 1 | 8 | 165pb | 0.0214 |
| Montpellier<br>(France) | 3 | 4-3-2 |  |  |
| Sapporo Japan) | 1 | 10 |  |  |
| Tokyo (Japan) | 1 | 8 |  |  |
| Dayton (U.S.A) | 1 | 9 |  |  |
| Watsonville<br>(U.S.A) | 1 | 10 |  |  |
|  |  | Jheh-2 intron 3 |  |  |
|  | N haplotype | N | Alignment | Mean diversity |
| Paris (France) | 1 | 8 | 211pb | 0.0333 |
| Montpellier<br>(France) | 1 | 8 |  |  |
| Sapporo Japan) | 1 | 10 |  |  |
| Tokyo (Japan) | 2 | 5-4 |  |  |
| Dayton (U.S.A) | 2 | 4-2 |  |  |
| Watsonville<br>(U.S.A) | 1 | 10 |  |  |
|  |  | Jheh-3 intron 2 |  |  |
|  | N haplotype | N | Alignment | Mean diversity |
| Paris (France) | 1 | 10 | 118pb | 0.0209 |
| Montpellier<br>(France) | 2 | 7-3 |  |  |
| Sapporo Japan) | 1 | 9 |  |  |
| Tokyo (Japan) | 3 | 7-2-1 |  |  |
| Dayton (U.S.A) | 1 | 9 |  |  |
| Watsonville<br>(U.S.A) | 1 | 10 |  |  |

**Table S3.** Pairwise genetic distance between genotypes. Bold values represent genetic diversity within lines.

| **Jheh-1.1** | **Paris** | **Montpellier** | **Sapporo** | **Tokyo** | **Dayton** | **Watsonville** |
| --- | --- | --- | --- | --- | --- | --- |
| Paris | **0.0000** |  |  |  |  |  |
| Montpellier | 0.0104 | **0.0000** |  |  |  |  |
| Sapporo | 0.0104 | 0.0000 | **0.0000** |  |  |  |
| Tokyo | 0.0208 | 0.0104 | 0.0104 | **0.0000** |  |  |
| Dayton | 0.0105 | 0.0000 | 0.0000 | 0.0105 | **0.0000** |  |
| Watsonville | 0.0104 | 0.0000 | 0.0000 | 0.0104 | 0.0000 | **0.0792** |
| **Jheh-1.2** | **Paris** | **Montpellier** | **Sapporo** | **Tokyo** | **Dayton** | **Watsonville** |
| Paris | **0.0000** |  |  |  |  |  |
| Montpellier | 0.0084 | **0.0169** |  |  |  |  |
| Sapporo | 0.0000 | 0.0084 | **0.0000** |  |  |  |
| Tokyo | 0.0000 | 0.0084 | 0.0000 | **0.0000** |  |  |
| Dayton | 0.0000 | 0.0084 | 0.0000 | 0.0000 | **0.0000** |  |
| Watsonville | 0.0000 | 0.0084 | 0.0000 | 0.0000 | 0.0000 | **0.0000** |
| **Jheh-1.3** | **Paris** | **Montpellier** | **Sapporo** | **Tokyo** | **Dayton** | **Watsonville** |
| Paris | **0.0000** |  |  |  |  |  |
| Montpellier | 0.0079 | **0.0000** |  |  |  |  |
| Sapporo | 0.0157 | 0.0236 | **0.0396** |  |  |  |
| Tokyo | 0.0079 | 0.0000 | 0.0236 | **0.0000** |  |  |
| Dayton | 0.0238 | 0.0159 | 0.0379 | 0.0159 | **0.0000** |  |
| Watsonville | 0.0236 | 0.0157 | 0.0226 | 0.0157 | 0.0152 | **0.0198** |
| **Jheh-2.1** | **Paris** | **Montpellier** | **Sapporo** | **Tokyo** | **Dayton** | **Watsonville** |
| Paris | **0.0460** |  |  |  |  |  |
| Montpellier | 0.0502 | **0.0502** |  |  |  |  |
| Sapporo | 0.0546 | 0.0462 | **0.0546** |  |  |  |
| Tokyo | 0.0502 | 0.0167 | 0.0378 | **0.0324** |  |  |
| Dayton | 0.0254 | 0.0466 | 0.0511 | 0.0466 | **0.0449** |  |
| Watsonville | 0.0502 | 0.0084 | 0.0462 | 0.0084 | 0.0466 | **0.0466** |
| **Jheh-2.2** | **Paris** | **Montpellier** | **Sapporo** | **Tokyo** | **Dayton** | **Watsonville** |
| Paris | **0.0000** |  |  |  |  |  |
| Montpellier | 0.0121 | **0.0081** |  |  |  |  |
| Sapporo | 0.0303 | 0.0303 | **0.0000** |  |  |  |
| Tokyo | 0.0061 | 0.0061 | 0.0242 | **0.0000** |  |  |
| Dayton | 0.0364 | 0.0364 | 0.0242 | 0.0303 | **0.0000** |  |
| Watsonville | 0.0000 | 0.0121 | 0.0303 | 0.0061 | 0.0364 | **0.0000** |
| **Jheh-2.3** | **Paris** | **Montpellier** | **Sapporo** | **Tokyo** | **Dayton** | **Watsonville** |
| Paris | **0.0000** |  |  |  |  |  |
| Montpellier | 0.0240 | **0.0000** |  |  |  |  |
| Sapporo | 0.0144 | 0.0096 | **0.0000** |  |  |  |
| Tokyo | 0.0144 | 0.0096 | 0.0000 | **0.0144** |  |  |
| Dayton | 0.0144 | 0.0096 | 0.0000 | 0.0000 | **0.0096** |  |
| Watsonville | 0.0825 | 0.0874 | 0.0777 | 0.0777 | 0.0777 | **0.0000** |
| **Jheh-3.2** | **Paris** | **Montpellier** | **Sapporo** | **Tokyo** | **Dayton** | **Watsonville** |
| Paris | **0.0000** |  |  |  |  |  |
| Montpellier | 0.0339 | **0.0339** |  |  |  |  |
| Sapporo | 0.0254 | 0.0085 | **0.0000** |  |  |  |
| Tokyo | 0.0339 | 0.0169 | 0.0085 | **0.0113** |  |  |
| Dayton | 0.0000 | 0.0339 | 0.0254 | 0.0339 | **0.0000** |  |
| Watsonville | 0.0339 | 0.0169 | 0.0085 | 0.0000 | 0.0339 | **0.0000** |

**Table S4.** Summary of the ANOVA 2 by gene and sex.

| Gene | Sex | Estimate | Df | Sum Sq | Mean Sq | F value | Pr(>F) |
| --- | --- | --- | --- | --- | --- | --- | --- |
| *Jheh-1* | Females | Treatment | 1 | 0.7578741 | 0.7578741 | 5.4488993 | 0.0286769 |
|  |  | Genotype | 5 | 5.7726966 | 1.1545393 | 8.3008099 | 0.0001334 |
|  |  | Treatment:Genotype | 5 | 7.6349146 | 1.5269829 | 10.9785737 | 0.0000172 |
|  |  | Residuals | 23 | 3.1990137 | 0.1390876 | NA | NA |
|  | Males | Treatment | 1 | 0.0375774 | 0.0375774 | 0.1612375 | 0.6915729 |
|  |  | Genotype | 5 | 1.5806168 | 0.3161234 | 1.3564255 | 0.2754043 |
|  |  | Treatment:Genotype | 5 | 1.4780750 | 0.2956150 | 1.2684280 | 0.3096968 |
|  |  | Residuals | 24 | 5.5933488 | 0.2330562 | NA | NA |
| *Jheh-2* | Females | Treatment | 1 | 1.3696002 | 1.3696002 | 13.3644443 | 0.0013171 |
|  |  | Genotype | 5 | 2.7850846 | 0.5570169 | 5.4353244 | 0.0019155 |
|  |  | Treatment:Genotype | 5 | 5.2867104 | 1.0573421 | 10.3174554 | 0.0000276 |
|  |  | Residuals | 23 | 2.3570606 | 0.1024809 | NA | NA |
|  | Males | Treatment | 1 | 0.2389486 | 0.2389486 | 4.2568387 | 0.0500705 |
|  |  | Genotype | 5 | 0.6101519 | 0.1220304 | 2.1739557 | 0.0908517 |
|  |  | Treatment:Genotype | 5 | 0.4690600 | 0.0938120 | 1.6712490 | 0.1799009 |
|  |  | Residuals | 24 | 1.3471889 | 0.0561329 | NA | NA |
| *Jheh-3* | Females | Treatment | 1 | 2.8083330 | 2.8083330 | 21.6508485 | 0.0001105 |
|  |  | Genotype | 5 | 1.8020725 | 0.3604145 | 2.7786163 | 0.0417902 |
|  |  | Treatment:Genotype | 5 | 3.4117019 | 0.6823404 | 5.2605045 | 0.0022994 |
|  |  | Residuals | 23 | 2.9833315 | 0.1297101 | NA | NA |
|  | Males | Treatment | 1 | 2.5825122 | 2.5825122 | 13.5076975 | 0.0011917 |
|  |  | Genotype | 5 | 4.5987363 | 0.9197473 | 4.8106909 | 0.0034625 |
|  |  | Treatment:Genotype | 5 | 0.6865256 | 0.1373051 | 0.7181674 | 0.6161005 |
|  |  | Residuals | 24 | 4.5885164 | 0.1911882 | NA | NA |

**Table S5.**
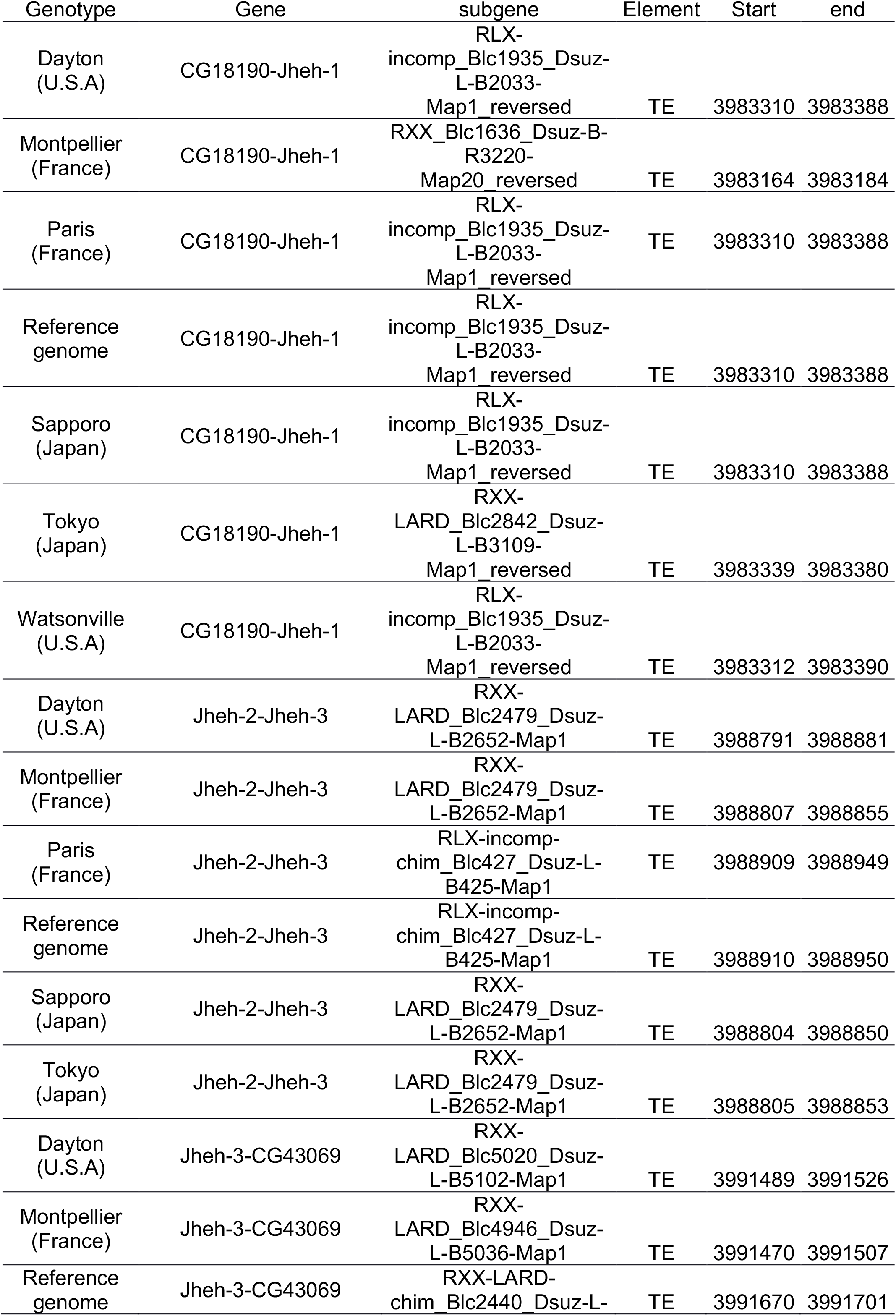
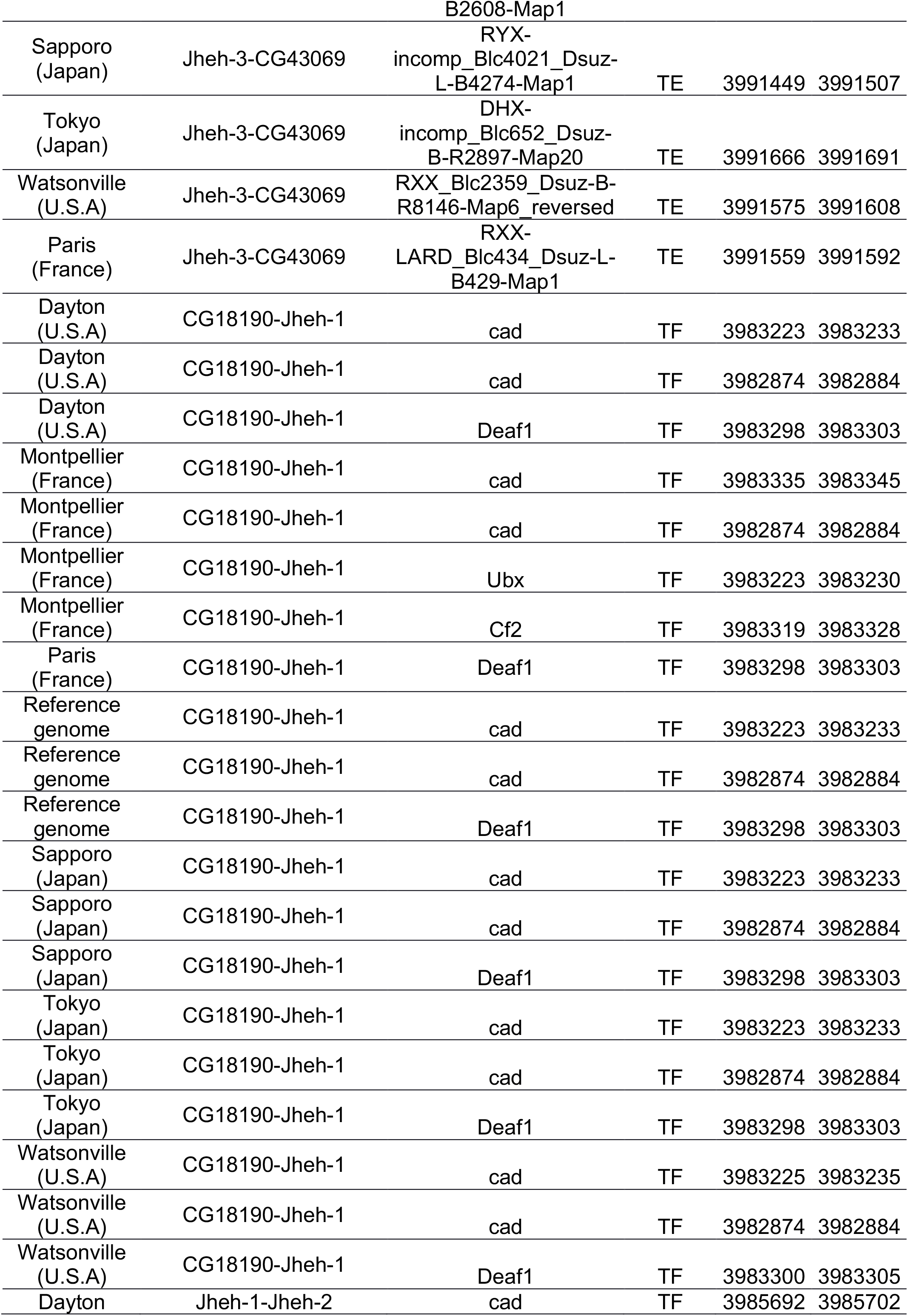
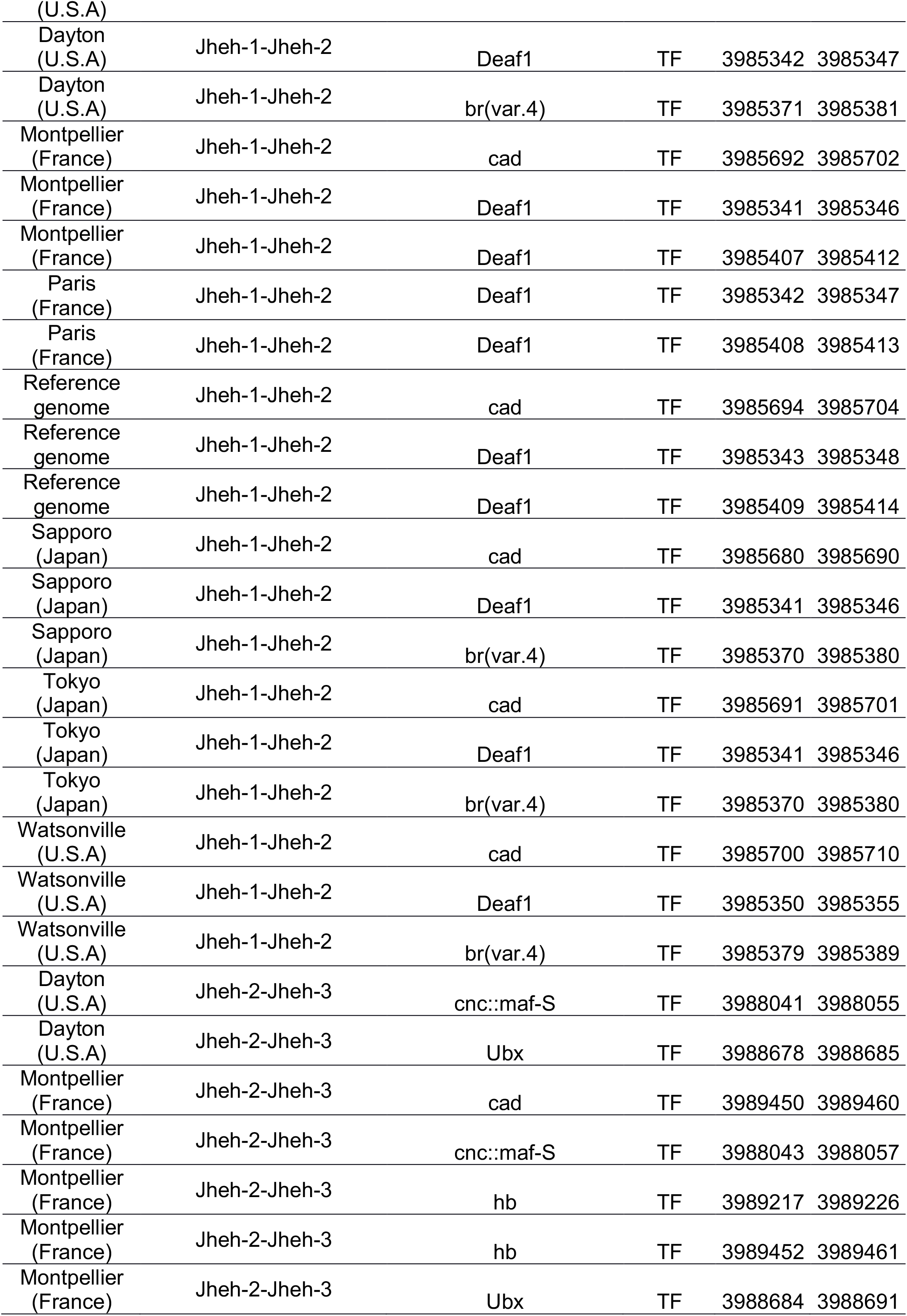
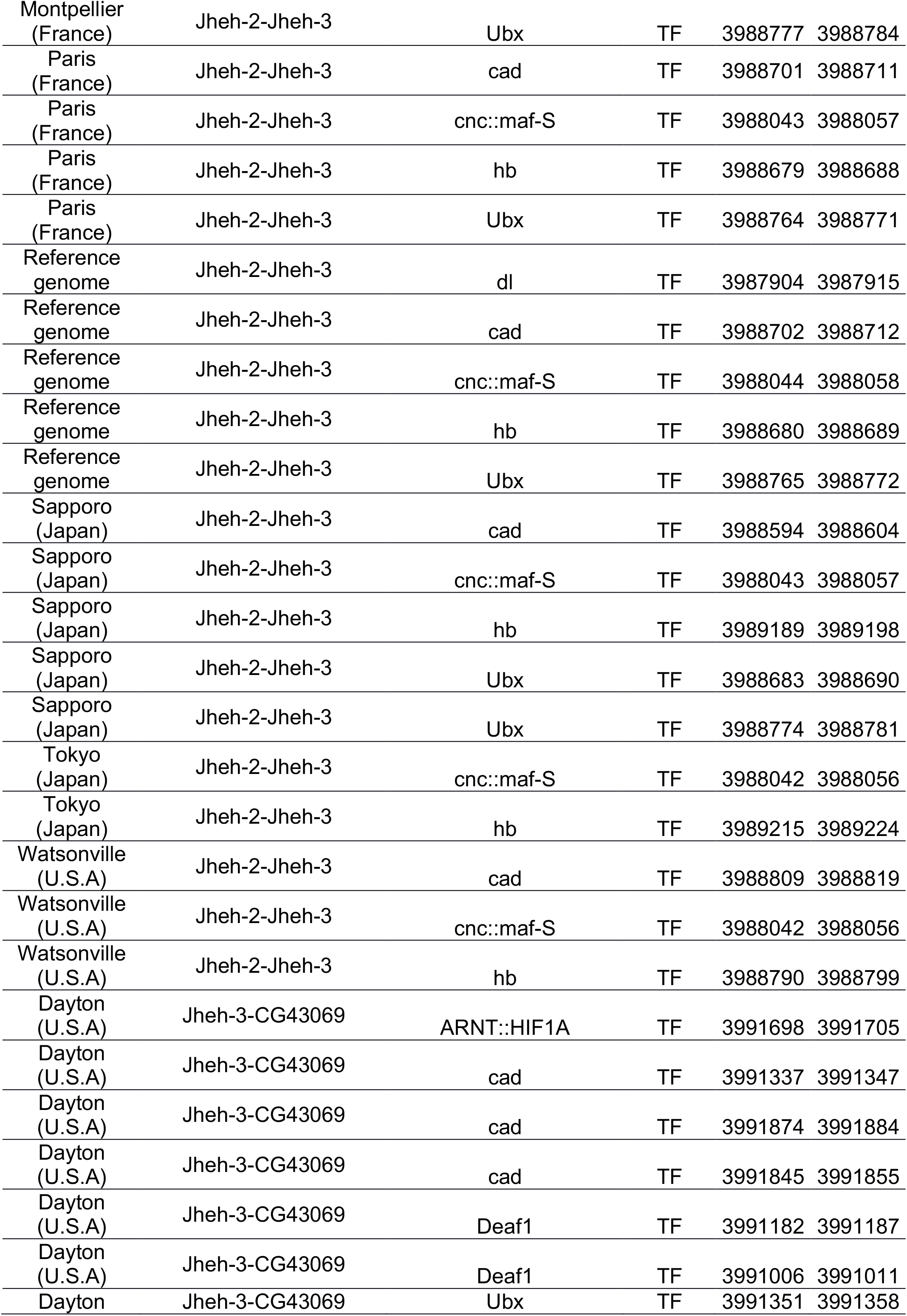
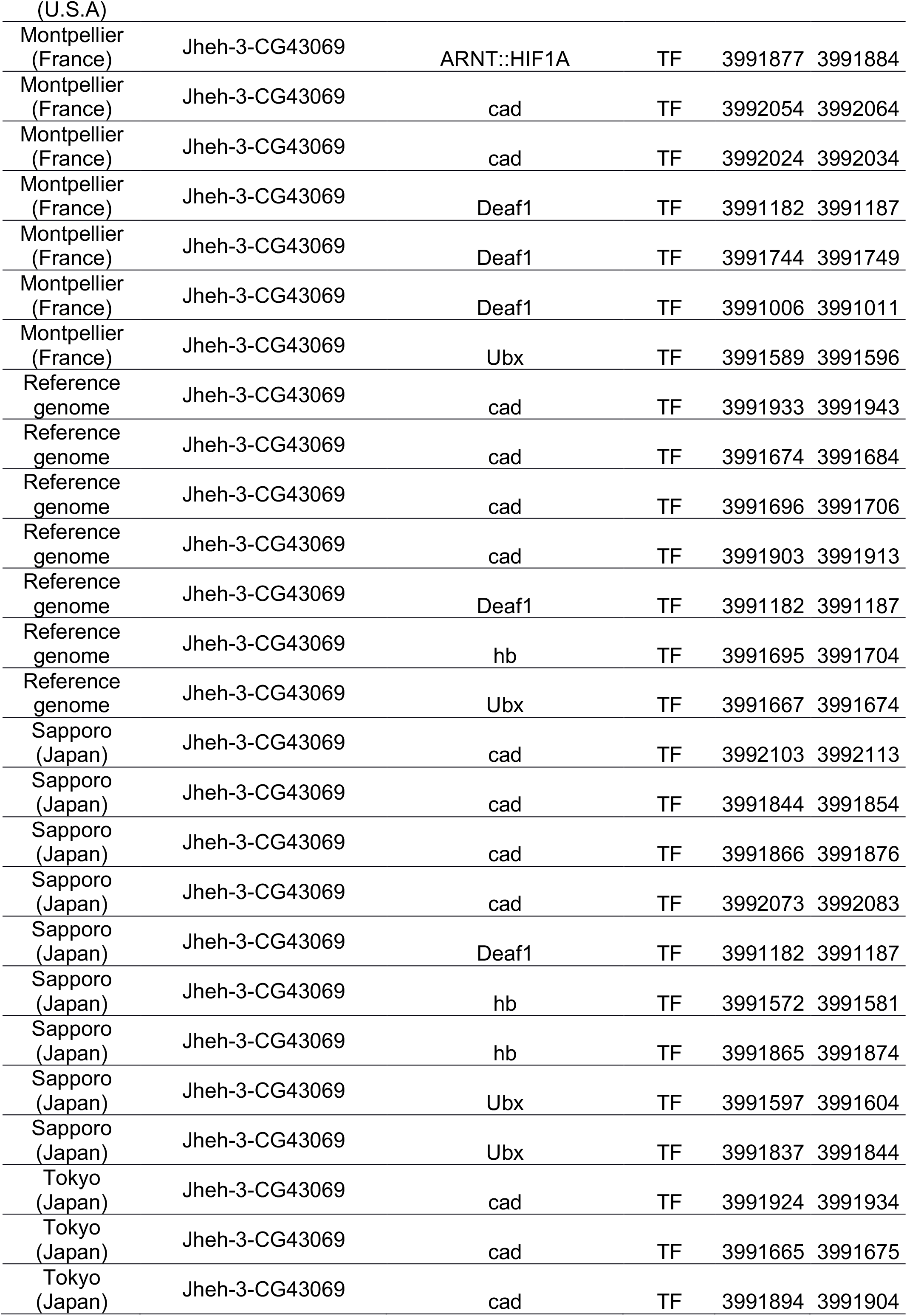
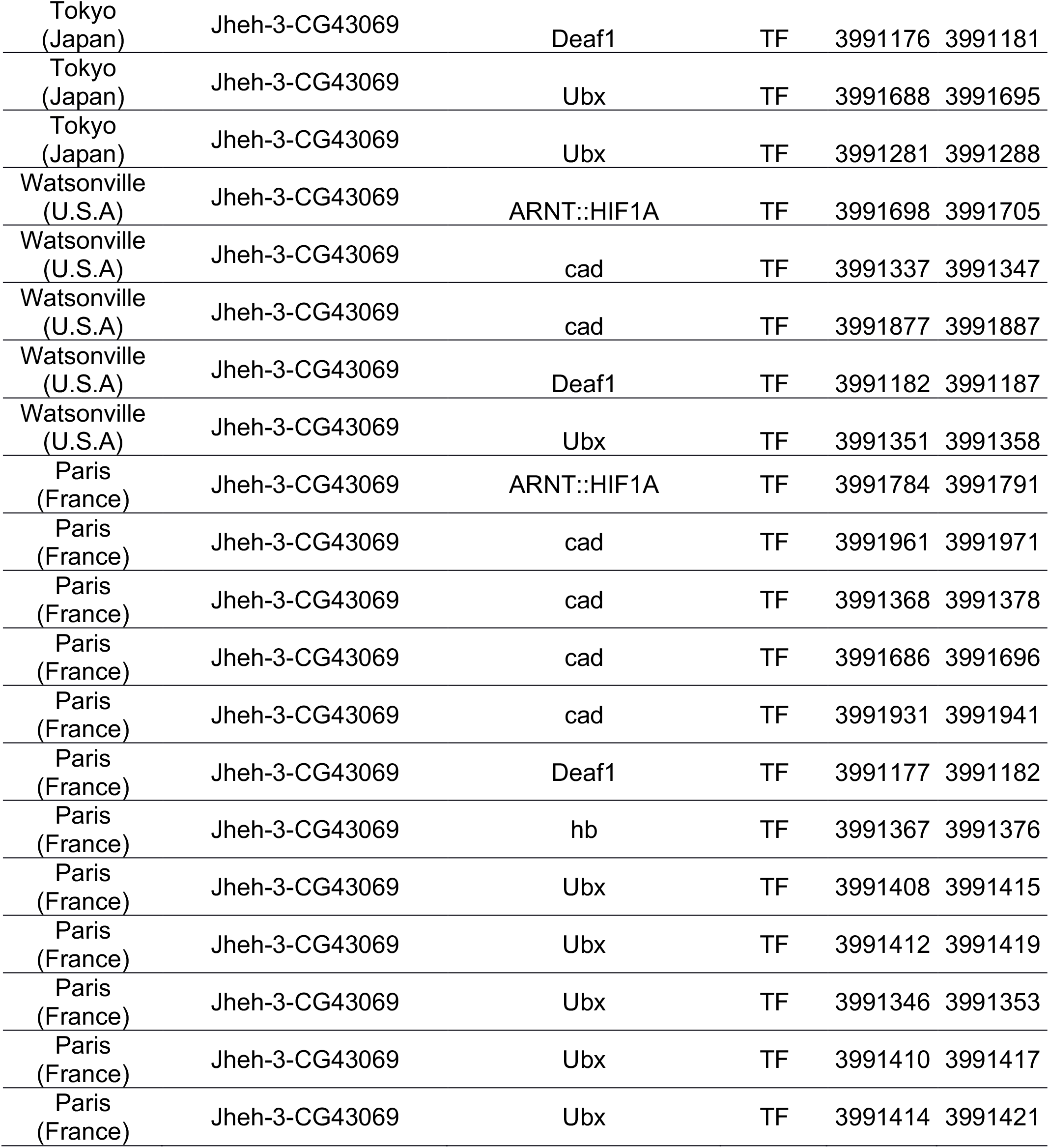
TFBS and TE detected in all six genotypes and the reference genome of *D. suzukii*. We screened TFBS in the intergenic regions before, between and after the Jheh genes. The names of the transcription factors (TF) and transposable elements (TE) are given with their positions in the sequence (beginning and end).

| Genotype | Gene | subgene | Element | Start | end |
| --- | --- | --- | --- | --- | --- |
| Dayton<br>(U.S.A) | CG18190-Jheh-1 | RLX-incomp_Blc1935_Dsuz-L-B2033-Map1_reversed | TE | 3983310 | 3983388 |
| Montpellier<br>(France) | CG18190-Jheh-1 | RXX_Blc1636_Dsuz-B-R3220-Map20_reversed | TE | 3983164 | 3983184 |
| Paris<br>(France) | CG18190-Jheh-1 | RLX-incomp_Blc1935_Dsuz-L-B2033-Map1_reversed | TE | 3983310 | 3983388 |
| Reference<br>genome | CG18190-Jheh-1 | RLX-incomp_Blc1935_Dsuz-L-B2033-Map1_reversed | TE | 3983310 | 3983388 |
| Sapporo<br>(Japan) | CG18190-Jheh-1 | RLX-incomp_Blc1935_Dsuz-L-B2033-Map1_reversed | TE | 3983310 | 3983388 |
| Tokyo<br>(Japan) | CG18190-Jheh-1 | RXX-LARD_Blc2842_Dsuz-L-B3109-Map1_reversed | TE | 3983339 | 3983380 |
| Watsonville<br>(U.S.A) | CG18190-Jheh-1 | RLX-incomp_Blc1935_Dsuz-L-B2033-Map1_reversed | TE | 3983312 | 3983390 |
| Dayton<br>(U.S.A) | Jheh-2-Jheh-3 | RXX-LARD_Blc2479_Dsuz-L-B2652-Map1 | TE | 3988791 | 3988881 |
| Montpellier<br>(France) | Jheh-2-Jheh-3 | RXX-LARD_Blc2479_Dsuz-L-B2652-Map1 | TE | 3988807 | 3988855 |
| Paris<br>(France) | Jheh-2-Jheh-3 | RLX-incomp-<br>chim_Blc427_Dsuz-L-B425-Map1 | TE | 3988909 | 3988949 |
| Reference<br>genome | Jheh-2-Jheh-3 | RLX-incomp-<br>chim_Blc427_Dsuz-L-B425-Map1 | TE | 3988910 | 3988950 |
| Sapporo<br>(Japan) | Jheh-2-Jheh-3 | RXX-LARD_Blc2479_Dsuz-L-B2652-Map1 | TE | 3988804 | 3988850 |
| Tokyo<br>(Japan) | Jheh-2-Jheh-3 | RXX-LARD_Blc2479_Dsuz-L-B2652-Map1 | TE | 3988805 | 3988853 |
| Dayton<br>(U.S.A) | Jheh-3-CG43069 | RXX-LARD_Blc5020_Dsuz-L-B5102-Map1 | TE | 3991489 | 3991526 |
| Montpellier<br>(France) | Jheh-3-CG43069 | RXX-LARD_Blc4946_Dsuz-L-B5036-Map1 | TE | 3991470 | 3991507 |
| Reference<br>genome | Jheh-3-CG43069 | RXX-LARD-<br>chim_Blc2440_Dsuz-L- | TE | 3991670 | 3991701 |

| B2608-Map1 |  |  |  |  |  |
| --- | --- | --- | --- | --- | --- |
| Sapporo<br>(Japan) | Jheh-3-CG43069 | RYX-incomp_Blc4021_Dsuz-L-B4274-Map1 | TE | 3991449 | 3991507 |
| Tokyo<br>(Japan) | Jheh-3-CG43069 | DHX-incomp_Blc652_Dsuz-B-R2897-Map20 | TE | 3991666 | 3991691 |
| Watsonville<br>(U.S.A) | Jheh-3-CG43069 | RXX_Blc2359_Dsuz-B-R8146-Map6_reversed | TE | 3991575 | 3991608 |
| Paris<br>(France) | Jheh-3-CG43069 | RXX-LARD_Blc434_Dsuz-L-B429-Map1 | TE | 3991559 | 3991592 |
| Dayton<br>(U.S.A) | CG18190-Jheh-1 | cad | TF | 3983223 | 3983233 |
| Dayton<br>(U.S.A) | CG18190-Jheh-1 | cad | TF | 3982874 | 3982884 |
| Dayton<br>(U.S.A) | CG18190-Jheh-1 | Deaf1 | TF | 3983298 | 3983303 |
| Montpellier<br>(France) | CG18190-Jheh-1 | cad | TF | 3983335 | 3983345 |
| Montpellier<br>(France) | CG18190-Jheh-1 | cad | TF | 3982874 | 3982884 |
| Montpellier<br>(France) | CG18190-Jheh-1 | Ubx | TF | 3983223 | 3983230 |
| Montpellier<br>(France) | CG18190-Jheh-1 | Cf2 | TF | 3983319 | 3983328 |
| Paris<br>(France) | CG18190-Jheh-1 | Deaf1 | TF | 3983298 | 3983303 |
| Reference<br>genome | CG18190-Jheh-1 | cad | TF | 3983223 | 3983233 |
| Reference<br>genome | CG18190-Jheh-1 | cad | TF | 3982874 | 3982884 |
| Reference<br>genome | CG18190-Jheh-1 | Deaf1 | TF | 3983298 | 3983303 |
| Sapporo<br>(Japan) | CG18190-Jheh-1 | cad | TF | 3983223 | 3983233 |
| Sapporo<br>(Japan) | CG18190-Jheh-1 | cad | TF | 3982874 | 3982884 |
| Sapporo<br>(Japan) | CG18190-Jheh-1 | Deaf1 | TF | 3983298 | 3983303 |
| Tokyo<br>(Japan) | CG18190-Jheh-1 | cad | TF | 3983223 | 3983233 |
| Tokyo<br>(Japan) | CG18190-Jheh-1 | cad | TF | 3982874 | 3982884 |
| Tokyo<br>(Japan) | CG18190-Jheh-1 | Deaf1 | TF | 3983298 | 3983303 |
| Watsonville<br>(U.S.A) | CG18190-Jheh-1 | cad | TF | 3983225 | 3983235 |
| Watsonville<br>(U.S.A) | CG18190-Jheh-1 | cad | TF | 3982874 | 3982884 |
| Watsonville<br>(U.S.A) | CG18190-Jheh-1 | Deaf1 | TF | 3983300 | 3983305 |
| Dayton | Jheh-1-Jheh-2 | cad | TF | 3985692 | 3985702 |
| (U.S.A) |  |  |  |  |  |
| Dayton<br>(U.S.A) | Jheh-1-Jheh-2 | Deaf1 | TF | 3985342 | 3985347 |
| Dayton<br>(U.S.A) | Jheh-1-Jheh-2 | br(var.4) | TF | 3985371 | 3985381 |
| Montpellier<br>(France) | Jheh-1-Jheh-2 | cad | TF | 3985692 | 3985702 |
| Montpellier<br>(France) | Jheh-1-Jheh-2 | Deaf1 | TF | 3985341 | 3985346 |
| Montpellier<br>(France) | Jheh-1-Jheh-2 | Deaf1 | TF | 3985407 | 3985412 |
| Paris<br>(France) | Jheh-1-Jheh-2 | Deaf1 | TF | 3985342 | 3985347 |
| Paris<br>(France) | Jheh-1-Jheh-2 | Deaf1 | TF | 3985408 | 3985413 |
| Reference<br>genome | Jheh-1-Jheh-2 | cad | TF | 3985694 | 3985704 |
| Reference<br>genome | Jheh-1-Jheh-2 | Deaf1 | TF | 3985343 | 3985348 |
| Reference<br>genome | Jheh-1-Jheh-2 | Deaf1 | TF | 3985409 | 3985414 |
| Sapporo<br>(Japan) | Jheh-1-Jheh-2 | cad | TF | 3985680 | 3985690 |
| Sapporo<br>(Japan) | Jheh-1-Jheh-2 | Deaf1 | TF | 3985341 | 3985346 |
| Sapporo<br>(Japan) | Jheh-1-Jheh-2 | br(var.4) | TF | 3985370 | 3985380 |
| Tokyo<br>(Japan) | Jheh-1-Jheh-2 | cad | TF | 3985691 | 3985701 |
| Tokyo<br>(Japan) | Jheh-1-Jheh-2 | Deaf1 | TF | 3985341 | 3985346 |
| Tokyo<br>(Japan) | Jheh-1-Jheh-2 | br(var.4) | TF | 3985370 | 3985380 |
| Watsonville<br>(U.S.A) | Jheh-1-Jheh-2 | cad | TF | 3985700 | 3985710 |
| Watsonville<br>(U.S.A) | Jheh-1-Jheh-2 | Deaf1 | TF | 3985350 | 3985355 |
| Watsonville<br>(U.S.A) | Jheh-1-Jheh-2 | br(var.4) | TF | 3985379 | 3985389 |
| Dayton<br>(U.S.A) | Jheh-2-Jheh-3 | cnc::maf-S | TF | 3988041 | 3988055 |
| Dayton<br>(U.S.A) | Jheh-2-Jheh-3 | Ubx | TF | 3988678 | 3988685 |
| Montpellier<br>(France) | Jheh-2-Jheh-3 | cad | TF | 3989450 | 3989460 |
| Montpellier<br>(France) | Jheh-2-Jheh-3 | cnc::maf-S | TF | 3988043 | 3988057 |
| Montpellier<br>(France) | Jheh-2-Jheh-3 | hb | TF | 3989217 | 3989226 |
| Montpellier<br>(France) | Jheh-2-Jheh-3 | hb | TF | 3989452 | 3989461 |
| Montpellier<br>(France) | Jheh-2-Jheh-3 | Ubx | TF | 3988684 | 3988691 |
| Montpellier<br>(France) | Jheh-2-Jheh-3 | Ubx | TF | 3988777 | 3988784 |
| Paris<br>(France) | Jheh-2-Jheh-3 | cad | TF | 3988701 | 3988711 |
| Paris<br>(France) | Jheh-2-Jheh-3 | cnc::maf-S | TF | 3988043 | 3988057 |
| Paris<br>(France) | Jheh-2-Jheh-3 | hb | TF | 3988679 | 3988688 |
| Paris<br>(France) | Jheh-2-Jheh-3 | Ubx | TF | 3988764 | 3988771 |
| Reference<br>genome | Jheh-2-Jheh-3 | dl | TF | 3987904 | 3987915 |
| Reference<br>genome | Jheh-2-Jheh-3 | cad | TF | 3988702 | 3988712 |
| Reference<br>genome | Jheh-2-Jheh-3 | cnc::maf-S | TF | 3988044 | 3988058 |
| Reference<br>genome | Jheh-2-Jheh-3 | hb | TF | 3988680 | 3988689 |
| Reference<br>genome | Jheh-2-Jheh-3 | Ubx | TF | 3988765 | 3988772 |
| Sapporo<br>(Japan) | Jheh-2-Jheh-3 | cad | TF | 3988594 | 3988604 |
| Sapporo<br>(Japan) | Jheh-2-Jheh-3 | cnc::maf-S | TF | 3988043 | 3988057 |
| Sapporo<br>(Japan) | Jheh-2-Jheh-3 | hb | TF | 3989189 | 3989198 |
| Sapporo<br>(Japan) | Jheh-2-Jheh-3 | Ubx | TF | 3988683 | 3988690 |
| Sapporo<br>(Japan) | Jheh-2-Jheh-3 | Ubx | TF | 3988774 | 3988781 |
| Tokyo<br>(Japan) | Jheh-2-Jheh-3 | cnc::maf-S | TF | 3988042 | 3988056 |
| Tokyo<br>(Japan) | Jheh-2-Jheh-3 | hb | TF | 3989215 | 3989224 |
| Watsonville<br>(U.S.A) | Jheh-2-Jheh-3 | cad | TF | 3988809 | 3988819 |
| Watsonville<br>(U.S.A) | Jheh-2-Jheh-3 | cnc::maf-S | TF | 3988042 | 3988056 |
| Watsonville<br>(U.S.A) | Jheh-2-Jheh-3 | hb | TF | 3988790 | 3988799 |
| Dayton<br>(U.S.A) | Jheh-3-CG43069 | ARNT::HIF1A | TF | 3991698 | 3991705 |
| Dayton<br>(U.S.A) | Jheh-3-CG43069 | cad | TF | 3991337 | 3991347 |
| Dayton<br>(U.S.A) | Jheh-3-CG43069 | cad | TF | 3991874 | 3991884 |
| Dayton<br>(U.S.A) | Jheh-3-CG43069 | cad | TF | 3991845 | 3991855 |
| Dayton<br>(U.S.A) | Jheh-3-CG43069 | Deaf1 | TF | 3991182 | 3991187 |
| Dayton<br>(U.S.A) | Jheh-3-CG43069 | Deaf1 | TF | 3991006 | 3991011 |
| Dayton | Jheh-3-CG43069 | Ubx | TF | 3991351 | 3991358 |
| (U.S.A) |  |  |  |  |  |
| Montpellier<br>(France) | Jheh-3-CG43069 | ARNT::HIF1A | TF | 3991877 | 3991884 |
| Montpellier<br>(France) | Jheh-3-CG43069 | cad | TF | 3992054 | 3992064 |
| Montpellier<br>(France) | Jheh-3-CG43069 | cad | TF | 3992024 | 3992034 |
| Montpellier<br>(France) | Jheh-3-CG43069 | Deaf1 | TF | 3991182 | 3991187 |
| Montpellier<br>(France) | Jheh-3-CG43069 | Deaf1 | TF | 3991744 | 3991749 |
| Montpellier<br>(France) | Jheh-3-CG43069 | Deaf1 | TF | 3991006 | 3991011 |
| Montpellier<br>(France) | Jheh-3-CG43069 | Ubx | TF | 3991589 | 3991596 |
| Reference<br>genome | Jheh-3-CG43069 | cad | TF | 3991933 | 3991943 |
| Reference<br>genome | Jheh-3-CG43069 | cad | TF | 3991674 | 3991684 |
| Reference<br>genome | Jheh-3-CG43069 | cad | TF | 3991696 | 3991706 |
| Reference<br>genome | Jheh-3-CG43069 | cad | TF | 3991903 | 3991913 |
| Reference<br>genome | Jheh-3-CG43069 | Deaf1 | TF | 3991182 | 3991187 |
| Reference<br>genome | Jheh-3-CG43069 | hb | TF | 3991695 | 3991704 |
| Reference<br>genome | Jheh-3-CG43069 | Ubx | TF | 3991667 | 3991674 |
| Sapporo<br>(Japan) | Jheh-3-CG43069 | cad | TF | 3992103 | 3992113 |
| Sapporo<br>(Japan) | Jheh-3-CG43069 | cad | TF | 3991844 | 3991854 |
| Sapporo<br>(Japan) | Jheh-3-CG43069 | cad | TF | 3991866 | 3991876 |
| Sapporo<br>(Japan) | Jheh-3-CG43069 | cad | TF | 3992073 | 3992083 |
| Sapporo<br>(Japan) | Jheh-3-CG43069 | Deaf1 | TF | 3991182 | 3991187 |
| Sapporo<br>(Japan) | Jheh-3-CG43069 | hb | TF | 3991572 | 3991581 |
| Sapporo<br>(Japan) | Jheh-3-CG43069 | hb | TF | 3991865 | 3991874 |
| Sapporo<br>(Japan) | Jheh-3-CG43069 | Ubx | TF | 3991597 | 3991604 |
| Sapporo<br>(Japan) | Jheh-3-CG43069 | Ubx | TF | 3991837 | 3991844 |
| Tokyo<br>(Japan) | Jheh-3-CG43069 | cad | TF | 3991924 | 3991934 |
| Tokyo<br>(Japan) | Jheh-3-CG43069 | cad | TF | 3991665 | 3991675 |
| Tokyo<br>(Japan) | Jheh-3-CG43069 | cad | TF | 3991894 | 3991904 |
| Tokyo<br>(Japan) | Jheh-3-CG43069 | Deaf1 | TF | 3991176 | 3991181 |
| Tokyo<br>(Japan) | Jheh-3-CG43069 | Ubx | TF | 3991688 | 3991695 |
| Tokyo<br>(Japan) | Jheh-3-CG43069 | Ubx | TF | 3991281 | 3991288 |
| Watsonville<br>(U.S.A) | Jheh-3-CG43069 | ARNT::HIF1A | TF | 3991698 | 3991705 |
| Watsonville<br>(U.S.A) | Jheh-3-CG43069 | cad | TF | 3991337 | 3991347 |
| Watsonville<br>(U.S.A) | Jheh-3-CG43069 | cad | TF | 3991877 | 3991887 |
| Watsonville<br>(U.S.A) | Jheh-3-CG43069 | Deaf1 | TF | 3991182 | 3991187 |
| Watsonville<br>(U.S.A) | Jheh-3-CG43069 | Ubx | TF | 3991351 | 3991358 |
| Paris<br>(France) | Jheh-3-CG43069 | ARNT::HIF1A | TF | 3991784 | 3991791 |
| Paris<br>(France) | Jheh-3-CG43069 | cad | TF | 3991961 | 3991971 |
| Paris<br>(France) | Jheh-3-CG43069 | cad | TF | 3991368 | 3991378 |
| Paris<br>(France) | Jheh-3-CG43069 | cad | TF | 3991686 | 3991696 |
| Paris<br>(France) | Jheh-3-CG43069 | cad | TF | 3991931 | 3991941 |
| Paris<br>(France) | Jheh-3-CG43069 | Deaf1 | TF | 3991177 | 3991182 |
| Paris<br>(France) | Jheh-3-CG43069 | hb | TF | 3991367 | 3991376 |
| Paris<br>(France) | Jheh-3-CG43069 | Ubx | TF | 3991408 | 3991415 |
| Paris<br>(France) | Jheh-3-CG43069 | Ubx | TF | 3991412 | 3991419 |
| Paris<br>(France) | Jheh-3-CG43069 | Ubx | TF | 3991346 | 3991353 |
| Paris<br>(France) | Jheh-3-CG43069 | Ubx | TF | 3991410 | 3991417 |
| Paris<br>(France) | Jheh-3-CG43069 | Ubx | TF | 3991414 | 3991421 |

